# Evolution of sperm competition: Natural variation and genetic determinants of *Caenorhabditis elegans* sperm size

**DOI:** 10.1101/501486

**Authors:** Clotilde Gimond, Anne Vielle, Nuno Silva-Soares, Stefan Zdraljevic, Patrick T. McGrath, Erik C. Andersen, Christian Braendle

**Author notes:** these authors contributed equally (Clotilde Gimond, Anne Vielle).

## Abstract

Sperm morphology is critical for sperm competition and thus for reproductive fitness. In the male-hermaphrodite nematode *Caenorhabditis elegans*, sperm size is a key feature of sperm competitive ability. Yet despite extensive research, the molecular mechanisms regulating *C. elegans* sperm size and the genetic basis underlying its natural variation remain unknown. Examining 97 genetically distinct *C. elegans* strains, we observe significant heritable variation in male sperm size but genome-wide association mapping did not yield any QTL (Quantitative Trait Loci). While we confirm larger male sperm to consistently outcompete smaller hermaphrodite sperm, we find natural variation in male sperm size to poorly predict male fertility and competitive ability. In addition, although hermaphrodite sperm size also shows significant natural variation, male and hermaphrodite sperm size do not correlate, implying a sex-specific genetic regulation of sperm size. To elucidate the molecular basis of intraspecific sperm size variation, we focused on recently diverged laboratory strains, which evolved extreme sperm size differences. Using mutants and quantitative complementation tests, we demonstrate that variation in the gene *nurf-1* – previously shown to underlie the evolution of improved hermaphrodite reproduction – also explains the evolution of reduced male sperm size. This result illustrates how adaptive changes in *C. elegans* hermaphrodite function can cause the deterioration of a male-specific fitness trait due to a sexually antagonistic variant, representing an example of intralocus sexual conflict with resolution at the molecular level. Our results further provide first insights into the genetic determinants of *C. elegans* sperm size, pointing at an involvement of the NURF chromatin remodelling complex.

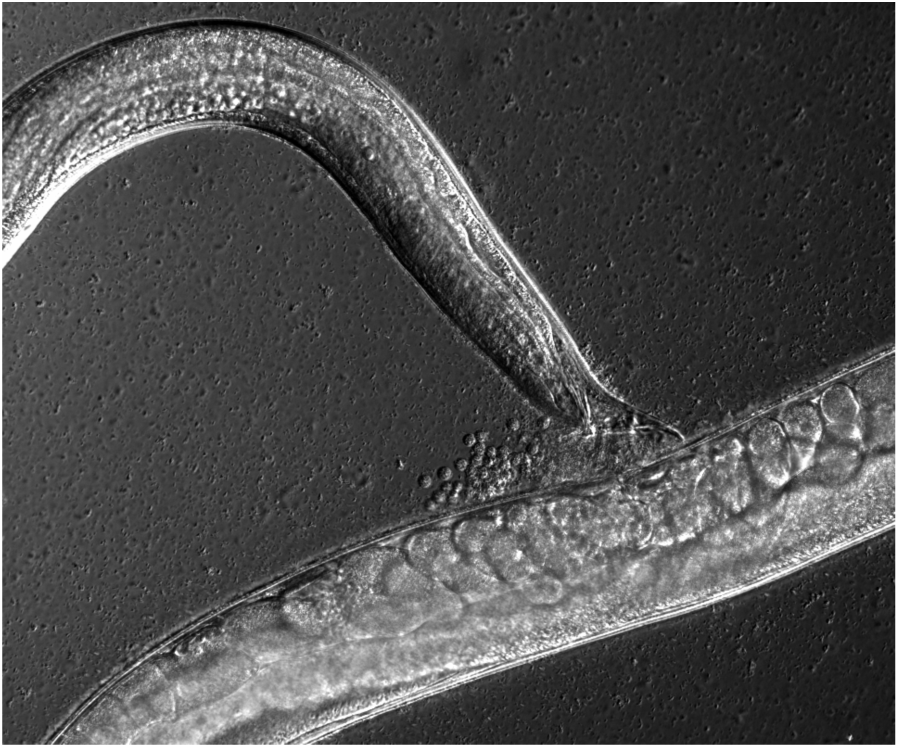

## INTRODUCTION

Sexual selection in the form of sperm competition is hypothesized to be responsible for rapid evolutionary divergence of sperm traits (Birkhead et al. 2009; Birkhead and Moller 1998; Pitnick et al. 2009; Ramm et al. 2014; Smith 1984; Snook 2005). In particular, sperm morphology may show extreme variation in size and shape among species, and is often associated with variation in competitive ability and thus male reproductive success. Furthermore, disparities of specific sperm traits, such as cell size or flagellum length, is not only common among species, but also within species (Pitnick et al. 2009). Numerous studies focusing on intraspecific variation, through comparison of sperm traits across populations of divergent individuals or by using artificial selection on sperm traits, have uncovered extensive levels of heritable variation in diverse sperm characteristics (Simmons and Moore 2009; Pitnick et al. 2009; Ward 1998; Joly et al. 2004; Morrow and Gage 2001a). Despite such studies on diverse invertebrate and vertebrate taxa, the quantitative and molecular genetic architecture of sperm traits associated with competitive ability remain largely undescribed. Therefore, although the developmental genetics of spermatogenesis and specific sperm traits have been elucidated in great detail from model organisms, such as *Drosophila melanogaster* (Demarco et al. 2014) or *Caenorhabditis elegans* (Ellis and Stanfield 2014), it is largely unknown whether uncovered genes are also a substrate for evolution to affect intraspecific variation in sperm characters relevant for male competitiveness.

Here we aimed to quantify and characterize intraspecific genetic variation of a well defined sperm trait (cell size) known directly to affect sperm competitive ability in the androdioecious (male-hermaphrodite) nematode *C. elegans*. Nematodes of the *Caenorhabditis* genus have become useful organisms to study sperm competition because it was demonstrated that the size of amoeboid *Caenorhabditis* sperm is a key factor for sperm competitive ability (Ward and Carrel 1979; LaMunyon and Ward 1995, 1998, 1999, 2002; Palopoli et al. 2015; Ting et al. 2014). In *C. elegans*, both males and hermaphrodites produce sperm, so that hermaphrodites can either self-fertilize or outcross with males; however, hermaphrodites cannot inseminate each other. *C. elegans* shows pronounced sperm size dimorphism: male sperm is larger and consistently outcompetes smaller hermaphrodite sperm when both types of sperm are present in the hermaphrodite reproductive tract (Ward and Carrel 1979; LaMunyon and Ward 1995). Male sperm size is further critical when multiple males compete for fertilization, with larger sperm outcompeting smaller sperm (LaMunyon and Ward 1998; Murray et al. 2011). In addition, increased levels of male-male competition in experimental contexts lead to the evolution of larger male sperm size (LaMunyon and Ward 2002; Palopoli et al. 2015), consistent with the relevance of sperm size for male competitive ability. Moreover, gonochoristic (male-female) *Caenorhabditis* species – with high levels of male-male competition – exhibit, on average, much larger male sperm than the three androdieocious species, *C. briggsae, C. elegans*, and *C. tropicalis*, in which male-male competition is much weaker (LaMunyon and Ward 1999; Vielle et al. 2016).

Although self-fertilization is the predominant mode of *C. elegans* reproduction, with reported rare occurrence of males and outcrossing events in natural populations (Jovelin et al. 2003; Barrière and Félix 2005; Sivasundar and Hey 2005), ample natural variation in diverse male traits exists (Hodgkin and Doniach 1997; Alcorn et al. 2016; Teotonio et al. 2006; Anderson et al. 2010; Noble et al. 2015; Morran et al. 2009; Palopoli et al. 2015; Palopoli et al. 2008), including male sperm size (LaMunyon and Ward 1998; Murray et al. 2011; Vielle et al. 2016). It remains unclear, however, to what degree such variation in male phenotypes reflects variation in male reproductive performance or competitive ability as a result of male-hermaphrodite or male-male interactions. Consistent with low rates of outcrossing, a number of male (and hermaphrodite) traits directly linked to mating, such as mate recognition, mating ability, or sex pheromone production are impaired, *e.g.* relative to related gonochoristic species (Sherlekar and Lints 2014; Kleemann and Basolo 2007; Noble et al. 2015; Lipton et al. 2004; Chasnov et al. 2007; Chasnov 2013; Garcia 2014; Thomas et al. 2012; Yin et al. 2018). In contrast, larger male sperm size was shown to clearly associate with increased male competitive ability and reproductive success, and male sperm size varies across *C. elegans* isolates (Ward and Carrel 1979; LaMunyon and Ward 1995, 1998, 1999, 2002; Palopoli et al. 2015; Murray et al. 2011). In the most comprehensive analysis to date, Murray *et al.* (2011) showed larger male sperm size to be a highly significant factor explaining natural variation in male fertilization success and male-male competitive ability across seven distinct *C. elegans* wild isolates. Extensive past research on *C. elegans* sperm competition thus indicates that male sperm size is tightly linked to male reproductive performance. Nevertheless, we have only limited knowledge of the extent of natural variation in *C. elegans* male sperm size, and we often ignore if this variation is adaptive as a result of (sexual) selection. Importantly, it also remains unclear how males and hermaphrodites generate differently sized sperm and to what extent sexual conflict in sperm size determination is resolved given that sperm size optima are strongly divergent between sexes: production of larger sperm will enhance male competitive ability while production of smaller sperm will accelerate hermaphrodite spermatogenesis to reduce developmental time until age of maturity, critical for reproductive fitness (Hodgkin and Barnes 1991; Cutter 2004). Currently, we lack information on how male sperm phenotypes, like other homologous traits shared across sexes, might be influenced by selection on hermaphrodite phenotypes given that hermaphrodite selfing is the predominant mode of reproduction in this species. In particular, we do not know whether male-hermaphrodite sperm size shows cross-sexual correlations across genotypes. Furthermore, although ample evidence exists for sperm traits such as length or size, these traits often co-vary with other traits, such as body size (allometry) or reproductive morphology of females (Pitnick et al. 2009).

Another large gap in our understanding of *C. elegans* sperm competition is the near-complete absence of information on the genetic determinants underlying sperm size variation and additional sperm phenotypes relevant for sperm competition. Although the genetic regulation of spermatogenesis has been elucidated in great detail (Chu and Shakes 2013; Ellis and Stanfield 2014; Geldziler et al. 2011; L’Hernault 2006), limited information is available on the molecular genetic control of sperm competitive ability (Hansen et al. 2015) and sperm size determination. No genes specifically affecting *C. elegans* sperm size have so far been identified. Spermatogenesis of *C. elegans* hermaphrodites and males seem essentially identical: meiosis is initiated by the formation of primary spermatocytes, followed by two rapid, mostly symmetrical divisions resulting in four haploid spermatids and an anucleate residual body (Shakes et al. 2009; Vielle et al. 2016; Ward et al. 1981). The cell size of the primary spermatocyte is a key determinant of final spermatid size (Vielle et al. 2016). Specifically, in *Caenorhabditis* nematodes, primary spermatocyte size differences between species, genotypes within species, and sexes within species strongly correlate with sperm size differences (Vielle et al. 2016). Therefore, sperm size seems to be mostly determined prior to or at the time of primary spermatocyte formation. However, it remains unknown how genetic factors contribute to this process. In addition, it is unclear to what extent the differential presence or activity of such potential genes explain reported differences in sperm size across sexes, genotypes, or species.

Here, we took advantage of a recently established world-wide collection of nearly 100 wild isolates (Andersen et al. 2012; Cook et al. 2017) to characterize the genetic basis of natural variation in *C. elegans* male sperm size. Our specific objectives were (i) to provide a quantitative assessment of natural variation in a presumed male-specific fitness trait in a partially selfing species; (ii) to perform a genome-wide association study to detect potential genetic loci explaining variation in male sperm size; (iii) to examine how observed natural variation in male sperm size correlates with variation in male reproductive performance and additional morphological traits of both males and hermaphrodites, including hermaphrodite sperm size; (iv) to characterize the molecular genetic basis of intraspecific variation in *C. elegans* male sperm.

## MATERIAL & METHODS

### Strains and culture conditions

All strains were maintained at 20°C on 2.5% agar NGM (Nematode Growth Medium) plates seeded with the *E. coli* strain OP50 (Stiernagle 2006). The following strains/genotypes were used in this study: 95 *C. elegans* wild isolates (Table S1) (Andersen et al. 2012; Cook et al. 2017) and laboratory strains N2, LSJ1, LSJ2, CX12311(*kyIR1* V, CB4856>N2; *qqIR1* X, CB4856>N2), CX13248 (*kyIR84* II, LSJ2>N2), *nurf-1(n4295)* (MT13649), *isw-1(n3294)* (MT17795), *isw-1(n3297)* (MT16012), *pyp-1(n4599)* IV/*nT1 [qIs51]* (MT14910), PD4790, and *C. plicata* (SB355). Hermaphrodites of the mutant *fog-2(q71)* (strain CB4108) do not produce any self-sperm, *i.e.* they are effectively females (Schedl and Kimble 1988), which were used for certain mating assays. The strain PD4790 contains an integrated transgene (*mls12* [*myo-2*::GFP, *pes-10*::GFP, *F22B7.9*::GFP]) in the N2 reference genetic background, expressing green fluorescent protein (GFP) in the pharynx. Additional information on wild isolates is available from CeNDR (http://elegansvariation.org) (Cook et al. 2017)

### Measurements of male and hermaphrodite sperm size

Males were collected from strain cultures at the L4 stage to be maintained on NGM plates containing only males to measure their spermatid size at stage L4+24h from synchronized and unmated males. Hermaphrodite spermatids were dissected from young virgin adults (at around mid L4+24h), at which stage most individuals contained both spermatids and activated sperm (spermatozoa), the latter of which were excluded from analysis. To measure sperm size, male or hermaphrodite spermatids were released into sperm medium (50mM HEPES pH7.8, 50mM NaCl, 25mM KCl, 5mM CaCl_2_, 1mM MgSO_4_, 1mg/ml BSA) by needle dissection (Nelson and Ward 1980). Images of the spermatids were captured using Nomarski optics (60x or 63x objectives). ImageJ software (Rasband, W.S., ImageJ, U. S. National Institutes of Health, Bethesda, Maryland, USA, http://imagej.nih.gov/ij/, 1997-2014) to calculate length and width of each spermatid to obtain measures of cross-sectional area assuming an ellipse shape: π x (length/2) x (width/2) (Vielle et al. 2016).

### Male mating ability

All mating assays (Figures 3 and 4) were performed on mating plates (35mm diameter NGM plates seeded with a spot of 20 µl OP50) using unmated males, *fog-2* females, or hermaphrodites that had been isolated at the L4 larval stage 24 hours (hermaphrodites and females) or 36 hours (males) prior to the assay. Following established protocols (Murray et al. 2011; Wegewitz et al. 2008), ten virgin *fog-2* females were transferred onto a mating plate and allowed to roam for 30 minutes on the bacterial lawn. Next, a single male was added to the plate and allowed to mate for eight hours, after which it was discarded. Females were left on the mating plate for an additional hour and were then picked as single animals onto fresh NGM plates. Females were scored as not fertilized if they did not contain any embryos in the uterus 24 hours later. Offspring production of fertilized females was followed over four consecutive days.

### Male sperm number transferred during single insemination (ejaculate size)

We quantified the number of sperm transferred during single insemination events (Figures 3E to 3H). For each strain to be tested (N2, CB4856, LSJ1, JU561, CX11285, EG4946, JU393, and JU782), unmated *fog-2* females were individually mated with an excess of 20-30 males, aged 36 hours post-L4 larval stage, to increase the chance of mating. Mating was monitored every few minutes by observation through the stereoscope. When a male was engaged in mating, it was kept under constant surveillance for spicule insertion and visualization of sperm flow from the male vas deferens to the female uterus, until mating was completed. Immediately after the end of mating, the inseminated female was isolated and fixed in ice-cold methanol and the mated male was removed from the male pool. A new virgin female was then mated with the males. Next, fixed females were washed twice in M9 and mounted in DAPI-containing Vectashield (Vector Laboratories, Inc., Burlingame, CA 94010). Sperm number was counted on images taken at 40x magnification as Z-stacks covering the entire thickness of the gonad using an Olympus BX61 microscope with a CoolSnap HQ2 camera (Poullet et al. 2016; Poullet et al. 2015).

### Hermaphrodite-male sperm competition

We measured variation in competition between hermaphrodite and male sperm by measuring male fertilization success of the eight strains when mated to hermaphrodites of a tester strain (the wild isolate CB4856) (Figure 4A). L4 hermaphrodites of the CB4856 strain were isolated 24 hours prior to mating, and L4 males of the eight strains were isolated 36 hours prior to mating. On the next day, one single CB4856 adult hermaphrodite was mated with an excess of 20-30 males for each strain to be tested and kept under surveillance for single mating as described above. As soon as mating was completed, the male was discarded from the male pool and the hermaphrodite was isolated onto a fresh NGM plate. Offspring (and male) production was scored for three days after the mating assay (*i.e.* until completion of the reproductive span).

### Male-male sperm competition

We measured second-male sperm precedence of the eight *C. elegans* strains using a tester strain expressing GFP in the pharynx (PD4790), following previously used protocols (Murray et al. 2011) (Figures 4C to 4E). *fog-2* females were first mated with PD4970 males, then with males of the eight strains. Males and *fog-2* females were isolated at the L4 stage and maintained in isolation for 36 hours prior to mating assays. Each mating plate (N=20) was established by adding ten *fog-2* females and 20 PD4790 males, which were allowed to mate for 15 hours, so that all females were fertilized (confirmed by the presence of embryos in the uterus). Ten Fertilized *fog-2* females were then randomly allocated to each new mating plate and allowed to mate with 20 males of each of the eight strains examined. Plates were kept under surveillance for single mating as described above. Upon completion of a mating event, both male and female were removed, and offspring production of the female was observed for the next four days. Total offspring were counted using a regular stereoscope and GFP-expressing offspring were counted with a fluorescence stereoscope.

### Quantification of hermaphrodite self-sperm number

We quantified the number of self-sperm in synchronized young adult hermaphrodites, *i.e.* adults containing one or two embryos in their uterus (Figure 5C). Animals were fixed overnight in ice-cold methanol (−20°C), washed three times in 1X PBS containing 0.05% Tween and mounted in Vectashield (Vector Lab) supplemented with DAPI. Sperm images were acquired from adults containing oocytes to ensure that the sperm to oocyte transition had occurred. Imaging of the anterior spermatheca was performed with an Olympus BX61 microscope using a 63X objective with epifluorescence. Z-sections (1 µm) of the entire spermatheca were taken and sperm number counted (Cell counter plugin in ImageJ) (Poullet et al. 2016; Poullet et al. 2015)

### Body size measurements

Synchronized populations were used to isolate unmated males in the mid-L4 stage and were scored 24 hours later. Hermaphrodites were scored as early adults when they contained between one and two embryos in the uterus. Animals were then anesthetized in sodium azide on an agar pad and whole-animal images were captured immediately after under Nomarski optics (20X). Body length and width were measured with the ImageJ software and body volume was calculated as that of a cylinder (π x (width/2)^2^ x length).

### Primary spermatocyte measurements

Extruded gonads from unmated males at 24 hours post-L4 were obtained by dissection in levamisole-containing M9. Gonads were fixed in 4% paraformaldehyde for 10 minutes and permeabilized for 5 minutes in 1X PBS 1X with 0.1% Triton X-100. Gonads were next stained for actin with phalloidin (1:500 dilution, Sigma-Aldrich) overnight at 4°C in a humidified chamber. Slides were mounted in Vectashield supplemented with DAPI (Vector Lab) and observed under an epifluorescent microscope. Primary spermatocyte area was measured by outlining cell boundaries using ImageJ software (Vielle et al. 2016).

### RNAi experiments

RNAi by bacterial feeding for *C. elegans* (N2) and *C. plicata* (SB355) was performed as previously described (Timmons and Fire 1998; Kamath et al. 2003). Briefly, control RNAi (HT115) and *nurf-1* clone (provided by the Ahringer lab) were seeded on standard NGM with 50ug/ml of ampicillin and 1mM of IPTG and grow at RT for at least 24h before experiment. Worms were fed RNAi and control bacteria from the L1 stage and spermatid size was measured in the early adult stage (L4+24h).

### Genome-wide association mapping

Genome-wide association (GWA) mapping was performed using phenotype data from 97 *C. elegans* isotypes (Table S2). We used the *cegwas* R package for association mapping (Cook et al. 2017). This package uses the EMMA algorithm for performing association mapping and correcting for population structure (Kang et al. 2008), which is implemented by the *GWAS* function in the *rrBLUP* package (Endelmann 2011). Specifically, the *GWAS* function in the *rrBLUP* package was called with the following command: *rrBLUP::GWAS(pheno = ph, geno = y, K = kin, min.MAF = 0.05, n.core = 1, P3D = FALSE, plot = FALSE)*. The kinship matrix used for association mapping was generated using a whole-genome high-quality single-nucleotide variant (SNV) set from CeNDR release 20160408 (Cook et al. 2016; Evans et al. 2017; Zdraljevic et al. 2017) and the *A.mat* function from the *rrBLUP* package. SNVs previously identified using RAD-seq (Andersen et al. 2012) that had at least 5% minor allele frequency in this strain set were used for performing GWA mappings. Burden test analyses were performed using RVtests (Zhan et al. 2016) and the variable-threshold method (Price et al. 2010). We called SNVs using bcftools (Li 2011) with settings previously described (Zhan et al. 2016; Cook et al. 2017; Cook et al. 2016). We next performed imputation using BEAGLE v4.1 (Cook et al. 2017) with *window* set to 8000, *overlap* set to 3000, and *ne* set to 17500. Within RVtests, we set the minor allele frequency range from 0.003 to 0.05 for burden testing.

### Statistical analyses

Statistical tests were performed using R, JMP, or SPSS. Data for parametric tests were transformed where necessary to meet the assumptions of ANOVA procedures (homogeneity of variances and normal distributions of residuals); all size data were log-transformed. For *post hoc* comparisons, Tukey’s honestly significant difference (HSD) procedure was used. For data, where ANOVA assumptions could not be met, we used nonparametric tests (*e.g.* Kruskal-Wallis).

Broad-sense heritability (*H*^2^) was estimated using the *lmer* function in the lme4 package (Bates et al. 2015) with the linear mixed model (phenotype ∼1 + (1|strain). *H*^2^ was then calculated as the fraction of the total variance explained by the random component (strain) of the mixed model.

### Data accessibility

All raw data are provided in Additional File S1.

## RESULTS

### Natural variation in *C. elegans* male sperm size

We quantified male sperm size variation of a world-wide collection of 97 *C. elegans* strains (Andersen et al. 2012), including two related laboratory strains (N2 and LSJ1), using measures of spermatid cross-sectional area. Average male sperm size, ranging from 15 µm^2^ to 27 µm^2^, varied significantly across strains (Figure 1 and Table S2). 90% of strains exhibited a male sperm size between 20 µm^2^ and 25 µm^2^, and we detected only two significant outliers: the wild strain JU561 (France) and the laboratory-adapted strain LSJ1 (McGrath et al. 2011) with the smallest male sperm size (Figures 1A and 1B). As found previously for *C. elegans* and other *Caenorhabditis* species (Vielle et al. 2016), we also detected high levels of inter-individual and intra-individual variation in male sperm size for most strains (Figures 1A and 1C).

**Figure 1.**
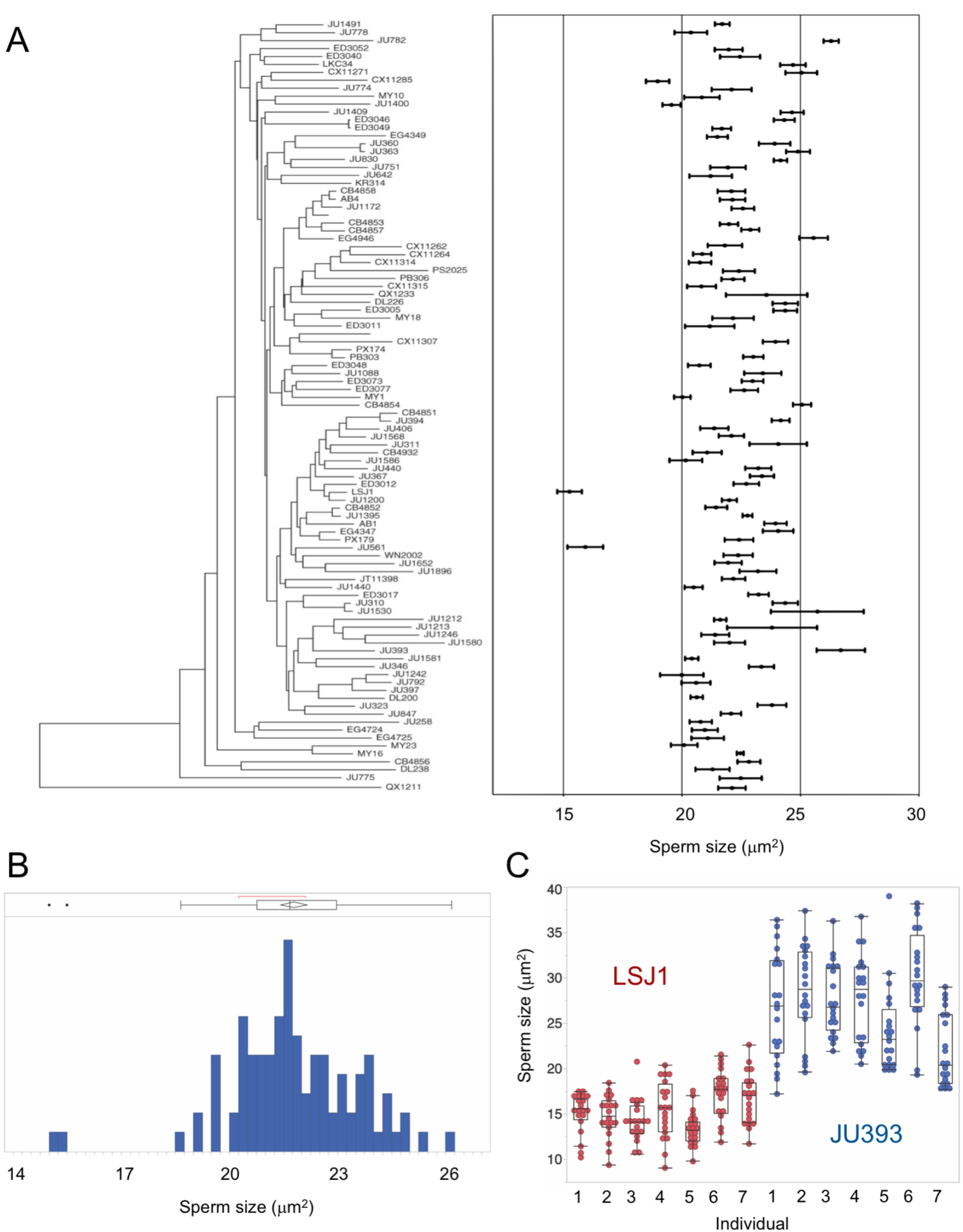
Natural variation in *C. elegans* male sperm size. (A) Quantification of male spermatid cross-sectional area area (mean ± sem) in 97 *C. elegans* strains, arranged with respect to the neighbour-joining tree of Andersen *et al.* (2012) based on genome-wide SNP data. There is significant genetic (and inter-individual) variation in sperm size (ANOVA, effect *strain*: F_96, 13539_ =24.75, P<0.0001; effect *individual(strain)*: F_580, 13539_ =3.08, P<0.0001). Twenty spermatids from each of seven individuals were measured per strain (N=140) with the exception of strains JU397 and KR413 for which 20 spermatids from each of six individuals (N=120) were measured (Table S2). (B) Histogram of sperm size across strains (least-squares mean estimates) shows the significant outlier trait values for the two strains (JU561 and LSJ1) with smallest male sperm. (C) Illustration of inter-and intra-individual variation in male sperm size for strains with smallest (LSJ1) versus largest male sperm size (JU393) (N=20 sperm per individual).

Coefficients of variation (CV), *i.e.* the ratio of the standard deviation to the mean, in sperm characters have been predicted – and shown – to be lower in species or genotypes experiencing higher levels of sperm competition (Fitzpatrick and Baer 2011; Gomendio et al. 2006; Immler et al. 2008; Calhim et al. 2007; Kleven et al. 2008). Therefore, we tested whether *C. elegans* strains with larger male sperm showed reduced variability. However, we did not detect a negative correlation between mean and CV of within-individual (ρ Pearson=0.14, P=0.17, N=97) or between-individual (ρ_Pearson_=-0.09, P=0.36, N=97) male sperm size (Table S2), as expected under such a scenario.

### Genome-wide association mapping of male sperm size

The significant variation in male sperm size enabled the mapping of genomic regions that could underlie this variation using genome-wide association studies, as has been performed successfully for a variety of traits using this *C. elegans* isolate panel (Andersen et al. 2012; Ashe et al. 2013; Ghosh et al. 2012). Broad-sense heritability was low (∼14%) for both sperm size cross-sectional area and diameter, and we found no significant genomic regions for these two traits and additional sperm size traits, including CV measurements (Figure 2). This result suggests that many loci could regulate differences in sperm size. Additionally, we used rare-variant based burden testing (Zhan et al. 2016; Bates et al. 2015; Price et al. 2010) to look for association of genes affected by deleterious rare variants with sperm area and diameter. As with marker-based association testing, we did not identify any significant genomic regions (data not shown).

**Figure 2.**
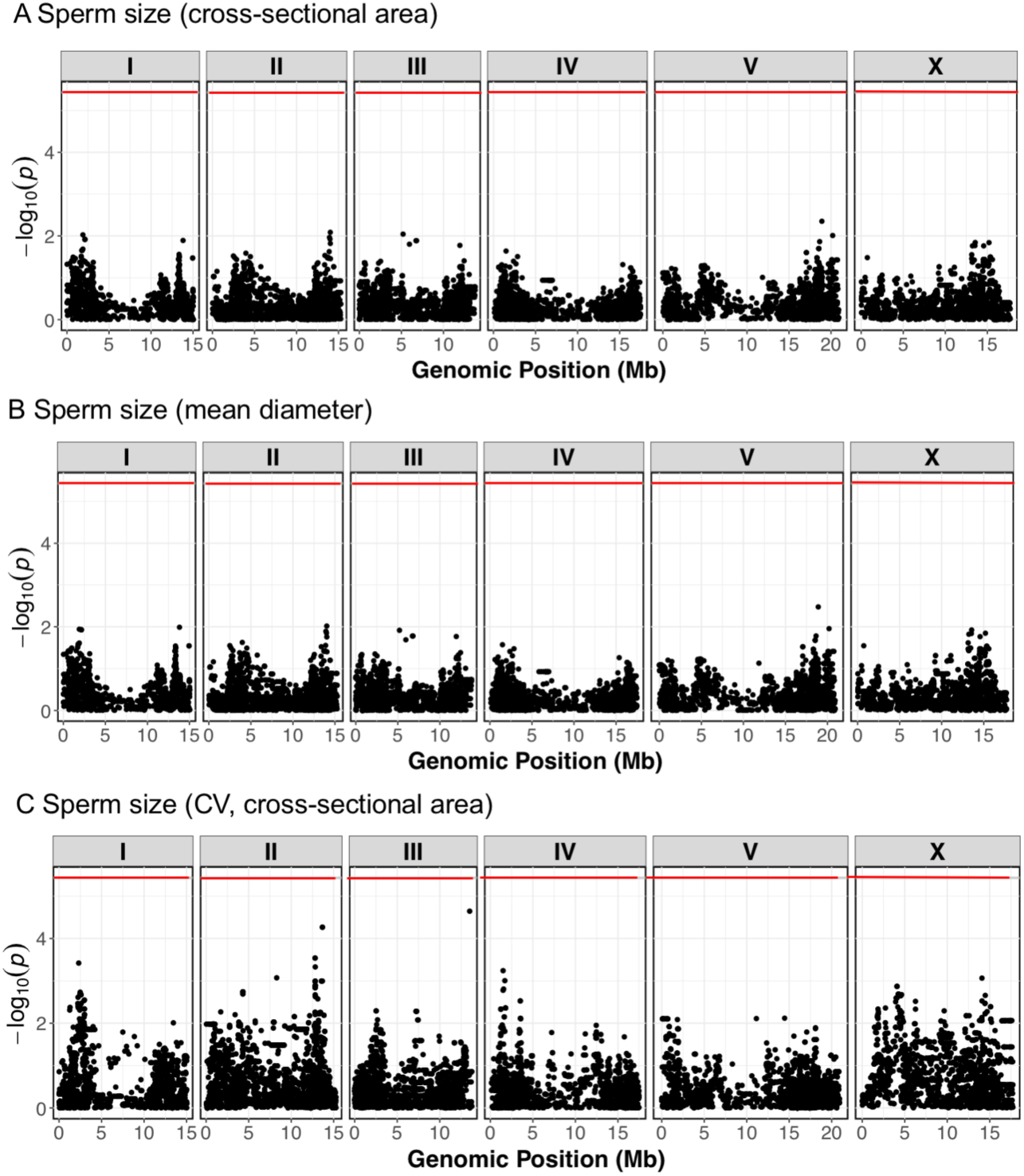
Genome-wide association mapping for *C. elegans* male sperm size. Manhattan plots of single-marker based GWA mappings show no significant genomic regions for least-squares mean estimates (LSM) of (A) sperm cross-sectional area, (B) sperm mean diameter and (C) CV (sperm cross-sectional area). Each dot represents an SNV that is present in at least 5% of the assayed population. The genomic location of each SNV is plotted on the x-axis, and the statistical significance is plotted on the y-axis. The Bonferroni-corrected significance threshold is shown as a red horizontal line.

Because specific natural niches could drive mating preferences, we investigated any effect of geography (*e.g.* latitude/longitude of strain origin) (Table S1, CeNDR: https://www.elegansvariation.org) on average male sperm size but found no such relationship (Spearman rank correlations, all P > 0.05).

### Sperm size is a poor predictor of male competitive ability and reproductive performance

Sperm size has been shown to be a central variable explaining differences in male reproductive success and competitive ability among *C. elegans* wild isolates (LaMunyon and Ward 1998; Murray et al. 2011; Wegewitz et al. 2008). We therefore tested whether the observed natural variation in *C. elegans* male sperm size correlates with variation in fertilization success and male competitive ability using eight strains with divergent male sperm size (from smallest to largest: LSJ1, JU561, CX11285, N2, CB4856, EG4946, JU782, JU393). Using similar assays described previously (Murray et al. 2011; Wegewitz et al. 2008), we measured the average number of inseminated *fog-2* females by a single male (Figure 3A) and resulting progeny counts (Figure 3C). For both measures, we found significant variation among strains (Figures 3A and 3C) but no correlation with sperm size (Figures 3B and 3D).

**Figure 3.**
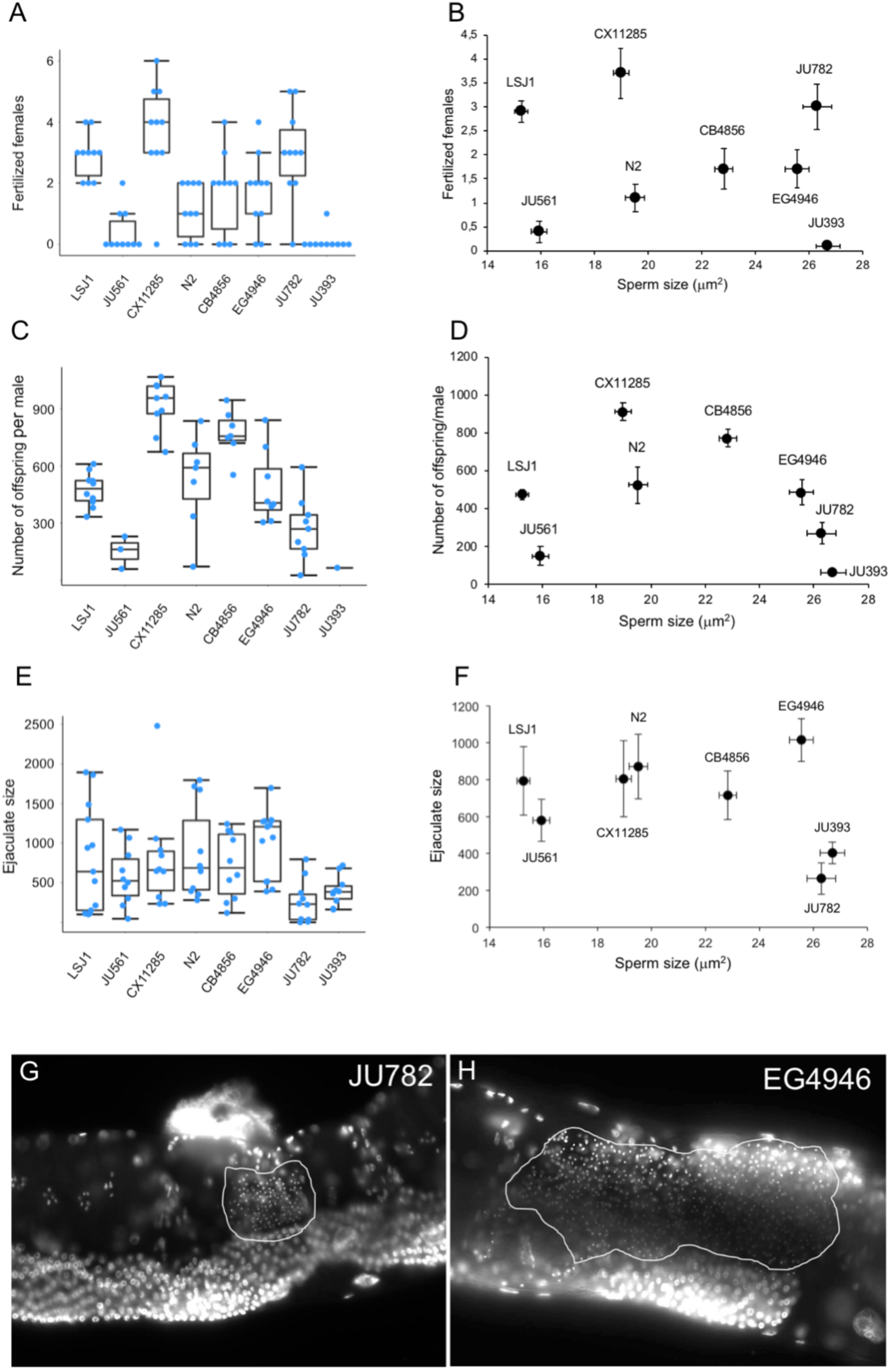
Differences in male mating ability and fertility between strains with variable male sperm size. Male mating ability, fertility, and ejaculate size in eight *C. elegans* strains with different average male sperm size. Strains with smallest (LSJ1) to largest (JU393) male sperm are arranged from left to right. (A) Significant strain variation in the number of *fog-2* females fertilized by a single male during an eight hour window (Kruskal-Wallis, χ^2^=45.12, df=7, P<0.0001) and (B) absence of correlation with average male sperm size (ρ_Spearman_=-0.23, P=0.59). (C) Significant strain variation in the number of offspring sired by a single male during eight hours of mating with up to ten females (Kruskal-Wallis, χ^2^=37.78, df=7, P<0.0001) and (D) absence of correlation with average male sperm size (ρ_Spearman_=-0.29, P=0.49). (E) Significant strain variation in ejaculate size, as measured by the number of sperm deposited by one male in a single mating (ANOVA, F_7, 86_=3.77, P=0.0014) and (F) no correlation between average ejaculate size and male sperm size (ρ_Spearman_=-0.33, P=0.42, log-transformed data). (G, H) Images of DAPI-stained *fog-2* females after single male mating event of strains (G) JU782 and (H) EG4946. Samples sizes were between 10 and 13 animals per strain per experiment.

We further measured ejaculate size, as the number of sperm present in the uterus and spermatheca of *fog-2* females after a single insemination event (Figure 3E). In contrast to a previous study on gonochoristic species *Caenorhabditis* species (Vielle et al. 2016), we did not find any evidence for a negative correlation between sperm size and ejaculate size (Figure 3F), which would be suggestive of a trade-off between sperm size and sperm number.

In addition, the polymorphic plugging phenotype of males, *i.e.* the deposition of a gelatinous plug on the vulva after copulation, likely representing a trait of male competitive ability as it affects mating ability of subsequent males (Palopoli et al. 2008; Hodgkin and Doniach 1997; Barker 1994), did not depend on sperm size as the average male sperm size did not differ between plugging versus non-plugging strains (ANOVA, F_1,96_ =0.09, P=0.77) (Table S3).

To complement the above experiments, we next asked whether variation in male sperm size may explain differences in male fertilization success when competing with (smaller) hermaphrodite self-sperm. In these experiments, young hermaphrodites of the strain CB4856 were allowed to mate once with a single male of the eight test strains. The proportion of resulting male progeny was then used as a measure of male fertilization success. Males of the eight strains showed significant differences in fertilization success when mated to CB4856 hermaphrodites (Figure 4A), however, there was no correlation between male fertilization success and sperm size across strains (Figure 4B). To test for differences in male-male competitive ability among the eight strains with different-sized sperm we used another, previously developed assay (Murray et al. 2011), in which *fog-2* females were first mated once to males of the transgenic strain PD4790, expressing green fluorescent protein (GFP) in the pharynx. Subsequently, these inseminated females were mated a second time, to a single male of the eight test strains. Competitive ability (in fertilization success) of a given wild strain is thus indicated by the extent of GFP-negative offspring. Although we found extensive variation in competitive ability (Figure 4C), no correlation between this measure of male sperm competitive ability and sperm size was observed (Figure 4D). Time-course analysis of sperm precedence over four consecutive days after mating revealed additional differences among strains, however, without any clear connection to differences in sperm size. (Figure 4E). Taken together, these experimental results suggest that *C. elegans* male sperm size does not consistently correlate with male fertilization success or competitive ability (Murray et al. 2011), implying that other sperm, morphological, or behavioural traits need to be considered to account for natural variation in male competitive ability and overall male reproductive performance. Consistent with this idea, we find that sperm transfer during a single mating (ejaculate size) (Figure 3E) rather than sperm size shows a strong positive correlation with male fertility when mated to hermaphrodites (Figure 4F).

**Figure 4.**
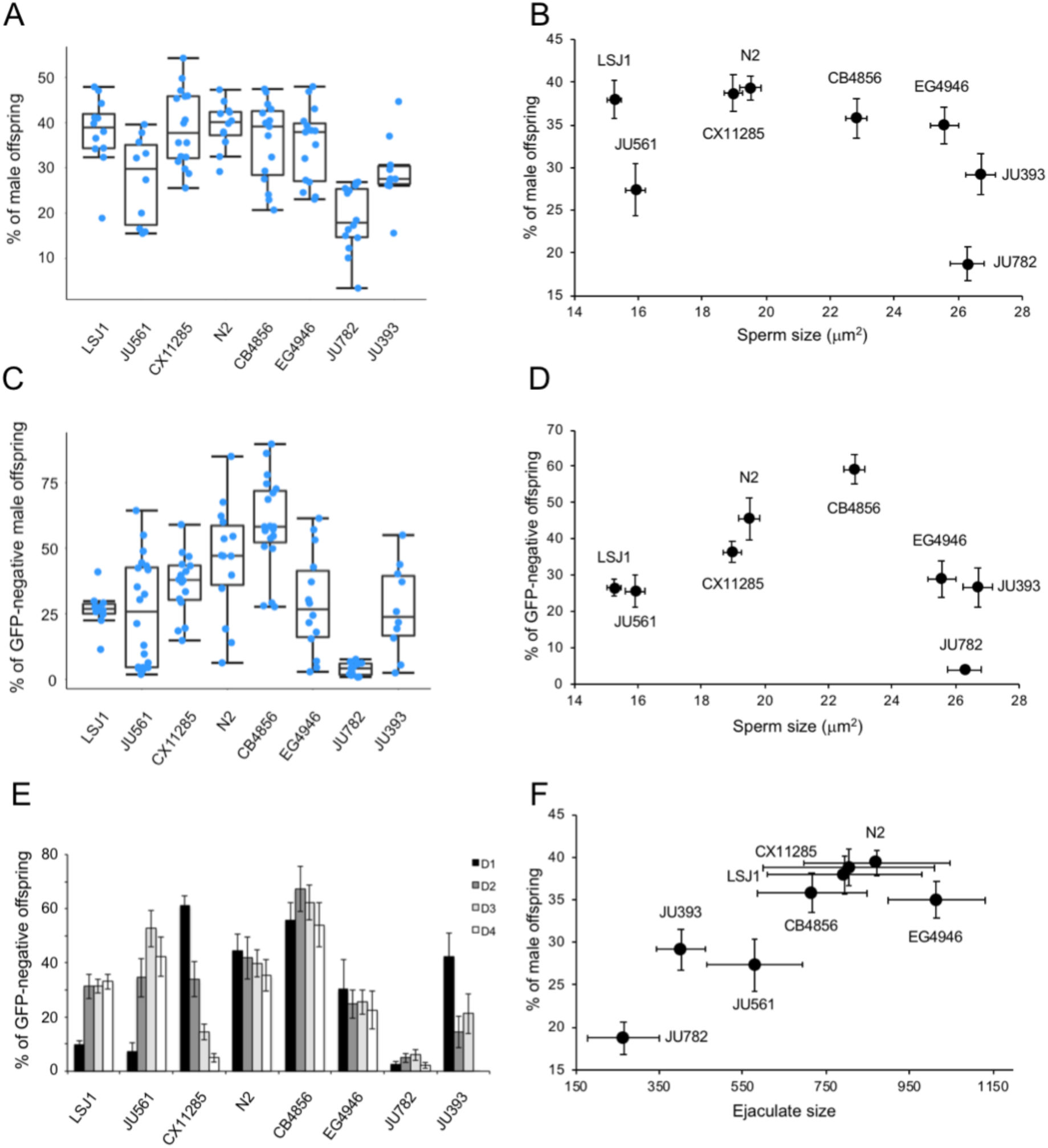
Differences in competitive ability between strains with variable male sperm size. Male competitive ability in fertilization success of eight *C. elegans* strains with different average male sperm size. Strains with smallest (LSJ1) to largest (JU393) sperm are arranged from left to right. (A) Significant strain variation in male fertilization success when competing with hermaphrodite self-sperm (strain CB4856) (ANOVA, F_7,103_=11.20, P<0.0001). (B) No correlation between male fertilization success and male sperm size (*ρ* _Spearman_=-0.45, P=0.45). (C) Significant strain variation in male-male competitive ability (in fertilization success) sperm (ANOVA, F_7,113_=14,66, P<0.0001). Male competitive ability of a given strain (versus the GFP-positive strain PD4790) quantified by the proportion of GFP-negative offspring produced over four days after mating. (D) No correlation between competitive ability and male sperm size (*ρ*_Spearman_=0.00, P=1). (E) Details of time course of progeny production across the four days after mating event (same data as in Figure 4C). (F) Significant correlation between ejaculate size (same data as in Figure 3E) and male fertilization success (Figure 4A) (*ρ*_Spearman_=0.74, P=0.036). Samples sizes: n=10-20 per strain per experiment.

### Natural variation in hermaphrodite sperm size: no correlation with variation in male sperm size

An unresolved question is whether genetic mechanisms in *C. elegans* sperm size determination are shared between the sexes, and more generally, to what extent homologous traits common to both sexes evolve. Hermaphrodite sperm is consistently smaller than male sperm in all three male-hermaphrodite species but also shows significant size variation across isolates within each species (Vielle et al. 2016). Previously, a weak positive correlation between the average sperm size of hermaphrodites and males was only found in *C. tropicalis* but not in *C. elegans* or *C. briggsae* (Vielle et al. 2016). This analysis was based on a small set of strains (n=5), so we revisited this question using 12 strains differing in male sperm size. In agreement with previous reports (Baldi et al. 2011; Vielle et al. 2016), hermaphrodite sperm showed significant genetic variation and was substantially smaller than male sperm in all strains (Figure 5A and Table S4). Again, as in Vielle et al. (2016), there was no significant cross-sexual correlation in sperm size (Figure 5B).

**Figure 5.**
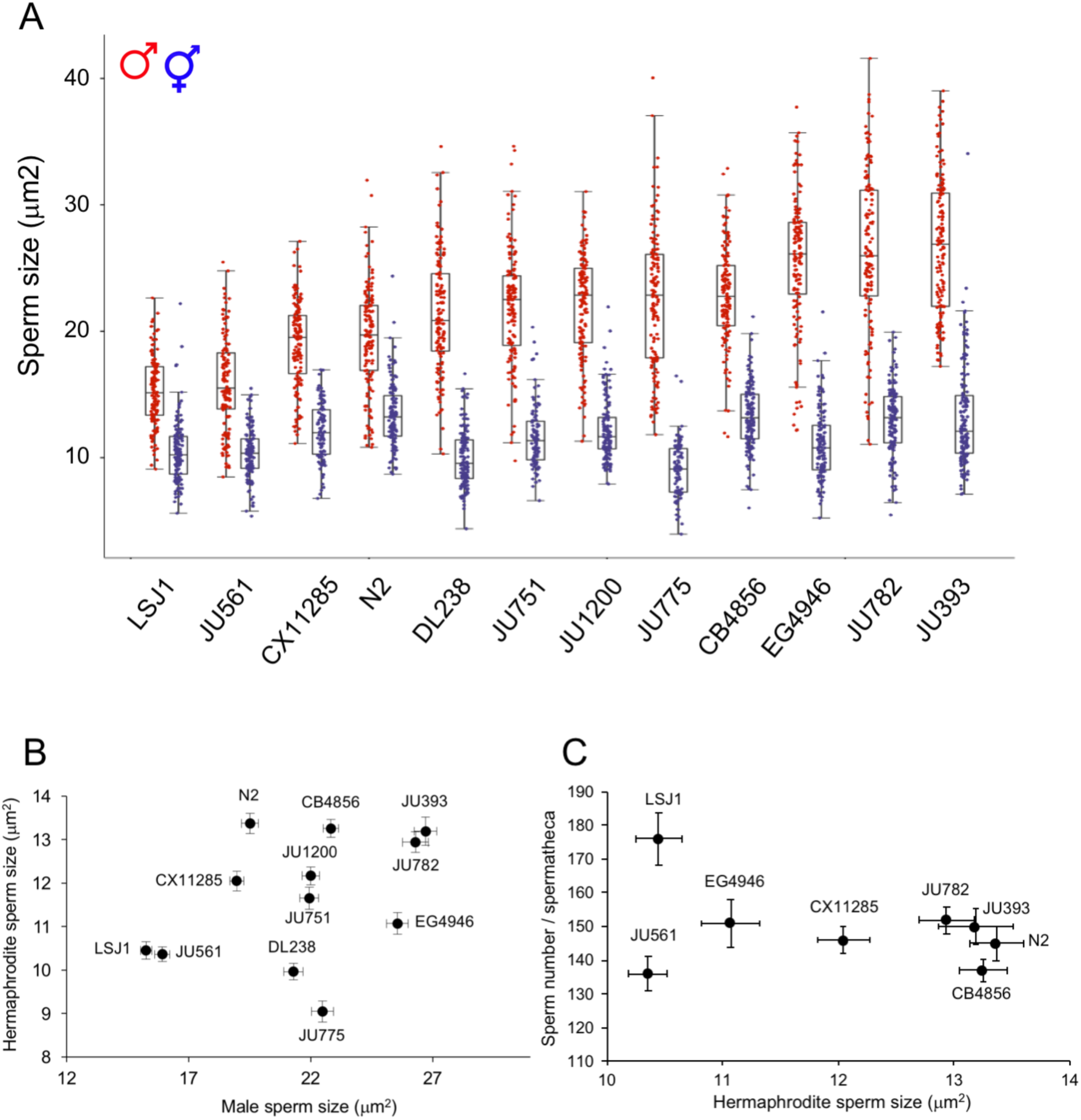
No correlation of *C. elegans* male and hermaphrodite sperm size across strains. (A) Effect of strain genotype and sex on *C. elegans* sperm size (ANOVA, effect *sex*: F_1,3207_=6267.55, P<0.0001; effect *strain:* F_11,3207_ =107.14, P<0.0001; interaction *sex* x *strain*: F_11, 3207_ =50.90, P<0.0001). For each strain, 87-152 hermaphrodite spermatids from 7-13 individuals were measured (male sperm size data from Figure 1). (B) The absence of a significant correlation between male and hermaphrodite sperm size was inferred from least-squares regression of strain mean values (F_1,12_=1.38, R^2^=0.11, P=0.27). (C) No correlation between average hermaphrodite sperm size and sperm number across eight strains (*ρ*_Pearson_=-0.039, P=0.40). Hermaphrodite self-sperm number were established by measuring all sperm contained within a single spermatheca of 12-31 individuals per strain.

Given the presence of significant natural variation in *C. elegans* hermaphrodite sperm size, we tested whether this variation is linked to differential sperm production, indicative of a potential trade-off between hermaphrodite sperm size and sperm number. Across eight strains hermaphrodite sperm production differed significantly (ANOVA, F_7,184_=5.37, P<0.0001) (Table S5), but we found no a negative correlation between hermaphrodite sperm size and number (Figure 5C).

Because sperm size and egg size show a weak positive correlation across (primarily gonochoristic) *Caenorhabditis* species (Vielle et al. 2016), we also tested for such a correlation across *C. elegans* strains. Using eggs size measurements from a previous study (Farhadifar et al. 2015), we did not detect any significant correlations, between either male sperm and egg size (F_1,72_=0.26, P=0.61) or between hermaphrodite sperm and egg size (F_1,9_=0.62, P=0.46).

### Allometric relationships between sperm and body size

We next tested whether natural variation in *C. elegans* sperm size may reflect fixed allometric relationships between body and cell size. Such positive correlations between sperm size (length or cell size) and animal body size or mass have been frequently observed in diverse invertebrate taxa (Pitnick et al. 2009). Size of amoeboid sperm of nematodes, including *Caenorhabditis*, partially correlates with male body size across species (LaMunyon and Ward 1999; Vielle et al. 2016). In contrast, whether intraspecific variation in body size is linked to variation in sperm size in *C. elegans*, and other species, had so far not been evaluated. Measuring early adult body size of hermaphrodites and males in a subset of strains, we found significant variation across strains and sex (Table S6); however, we did not detect any positive correlation between average sperm size and body size (length) in either sex (Figures 6A and 6B). In males only, sperm size was strongly correlated with body width (F_1,10_=16.86, R^2^=0.65, P=0.0027) (Figures 6C and 6D). As male body width may simply increase due to storing of larger sperm and thus increased gonad width (Vielle et al. 2016), we found little evidence for a positive scaling relationship between male body size and sperm size.

**Figure 6.**
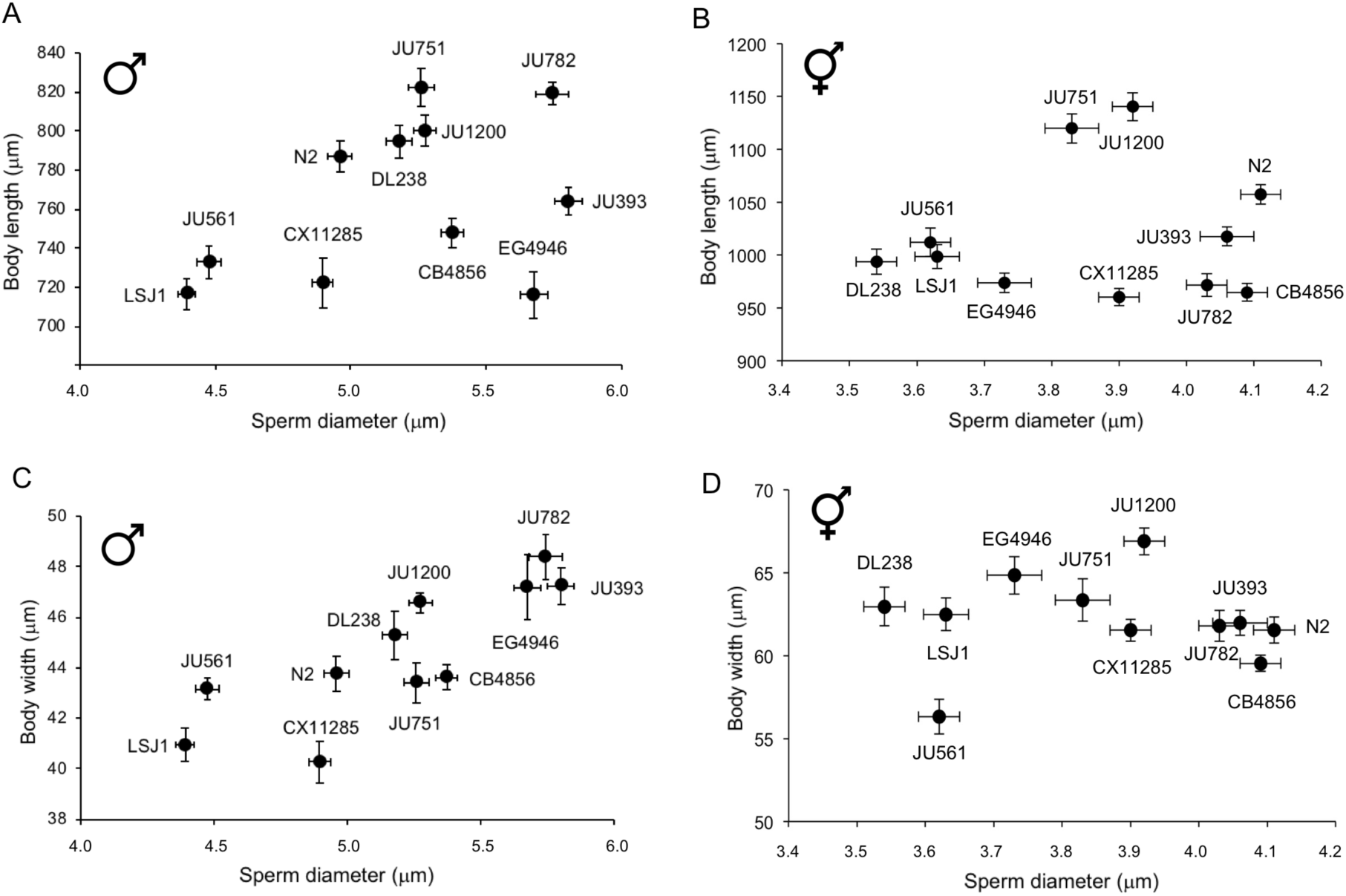
Allometric relationships between sperm and body size. Absence of scaling relationships between sperm size (mean diameter) and body length in (A) males (F_1,10_=1.88, R^2^=0.17, P=0.20) and (B) hermaphrodites (F_1,10_=0.005, R^2^=0.0005, P=0.94). Significant positive association between body width and sperm size in (C) males (F_1,10_=16.61, R^2^=0.65, P=0.0028) but not (D) hermaphrodites (F_1,10_=0.43, R^2^=0.05, P=0.53). Sperm size measures based on least-squares regression of strain mean (LSM); all data log-transformed for statistical analyses. Body length and width was measured in 10-29 individuals per strain and per sex (Table S6).

### Laboratory-derived strains LSJ1 and LSJ2 exhibit strongly reduced male and hermaphrodite sperm size

Across all 97 *C. elegans* strains measured, LSJ1 exhibited the smallest male sperm size (Figure 1A). LSJ1 is a laboratory-derived strain and shares a common ancestor with the reference strain N2 (Figure 7A), which shows a much larger male sperm size (Figures 1A and 7B). The two lineages diverged between 1957-1958. N2 was then maintained on agar plates seeded with *E. coli* for approximately 15 years, while LSJ1 was maintained in axenic liquid culture for close to 40 years before cultures were cryopreserved. LSJ2 is a derivative of LSJ1 that was kept in liquid culture for another 14 years prior to freezing (McGrath et al. 2011; McGrath et al. 2009; Large et al. 2017; Sterken et al. 2015). We therefore also measured LSJ2, which displayed a similarly small sperm size as LSJ1 (Figure 7B). In contrast, the N2-derived strain CC1, grown in liquid axenic medium for only four years, did not differ from N2 in male sperm size (Figure 7B). Given the common, inbred, and likely isogenic “Bristol ancestor” of LSJ and N2-CC1 lineages, these results suggest that the evolution of reduced male sperm size in the LSJ lineage occurred due to *de novo* mutations before 1995. Of note, LSJ1 and LSJ2 strain also showed significantly reduced hermaphrodite sperm size relative to N2 (and CC1) (Figure 7C).

**Figure 7.**
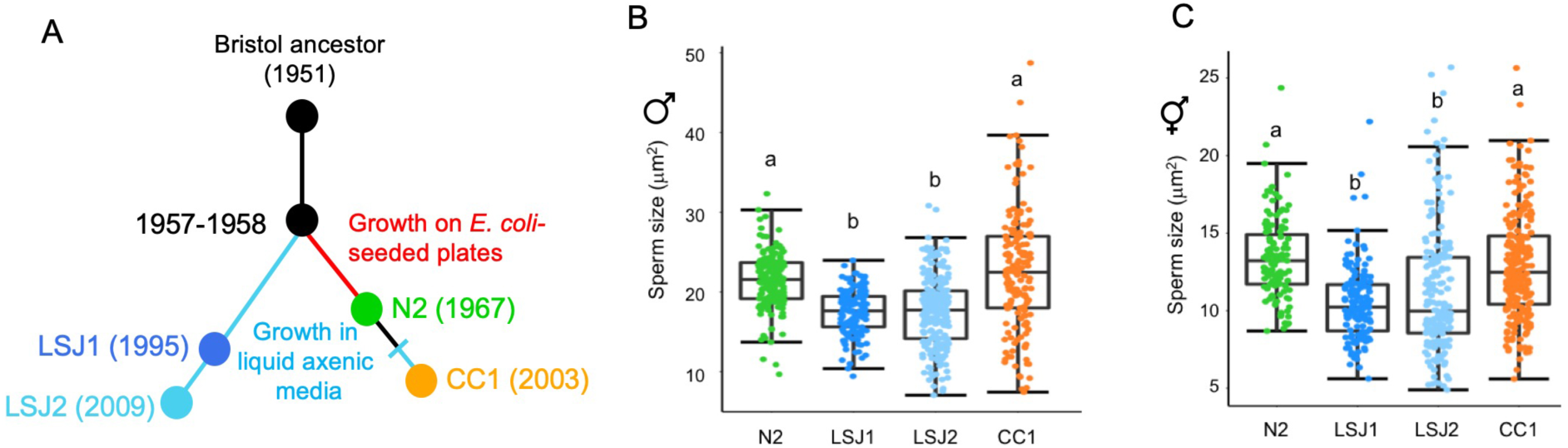
LSJ1 and LSJ2 strains exhibit strongly reduced male and hermaphrodite sperm size. (A) Laboratory evolution of LSJ and N2 lineages in the laboratory, after isolation of the common ancestral strain “Bristol”, derived from a single hermaphrodite individual, in 1951. LSJ1 and LSJ2 were cultivated in axenic liquid medium and N2 was cultivated on agar plates (after McGrath et al. 2011). B) Male sperm size: LSJ1 and LSJ2 exhibit significantly reduced male sperm size compared to N2 and CC1 (ANOVA, effect *strain*: F_3,658_=65.62, P<0.0001). (C) Hermaphrodite sperm size: LSJ1 and LSJ2 exhibit significantly reduced hermaphrodite sperm size compared to N2 and CC1 (ANOVA, effect *strain*: F_3,666_=40.95, P<0.0001). For male sperm measurements, 135-225 sperm were analysed from 9-15 individuals of each strain. For hermaphrodite sperm measurements, 123-248 sperm were analysed from 9-19 individuals of each strain. Values labelled with different letters indicate significant differences (Tukey’s HSD, *P*<0.05).

### Reduced male sperm size of LSJ strains is caused by genetic variation in *nurf-1*

Whole-genome short-read sequencing identified 188 and 94 new mutations fixed in the LSJ2 and N2 lineages, respectively (McGrath et al. 2011). These mutations include a 60 bp deletion in the 3’ end of *nurf-1*, which encodes the orthologue of the BPTF subunit of the NURF chromatin remodelling complex (McGrath et al. 2011; Large et al. 2016; Andersen et al. 2006). This variant is predicted to replace the last 16 amino acids of the protein with 11 novel residues and is known to underlie multiple life history differences between N2 and LSJ2, including reproductive timing, progeny production, growth rate, lifespan and dauer formation (Large et al. 2017; Large et al. 2016). Moreover, the NURF complex has previously been shown to function in *C. briggsae* spermatogenesis (Chen et al. 2014) as well as *Drosophila* spermatogenesis (Kwon et al. 2009). We thus reasoned that the *nurf-1* deletion specific to the LSJ lineage provides a good candidate explaining reduced male (and hermaphrodite) sperm size. This hypothesis was supported by the observation that RNAi knock-down of *nurf-1* resulted in a significant reduction of male sperm size in the N2 strain (Figure S1). To further test our hypothesis, we first examined the introgression line CX13248 (*kyIR87*) containing the LSJ2 region surrounding *nurf-1* in an N2-like background (CX12311, which is of N2 genotype except for introgressed *npr-1* and *glb-5* alleles from the strain CB4856) (McGrath et al. 2011). The *kyIR87* introgression contains the 60 bp deletion along with LSJ2 alleles of eight additional variants, including an SNV in the intron of *nurf-1* that was fixed in the N2 lineage (Large et al. 2016). Consistent with our hypothesis, sperm size of the CX13248 strain containing the *nurf-1* deletion was significantly smaller compared to CX12311, both in males (Figure 8A) and hermaphrodites (Figure 8B). In addition, sperm size of the *nurf-1(n4295)* deletion mutant (N2 background) (Andersen et al. 2006) was also strongly reduced in both sexes (Figures 8C and 8D); therefore, strains containing two different deletions in the 3’ coding region of *nurf-1* result in reduced *C. elegans* sperm size. Specifically, the 60 bp *nurf-1* deletion of LSJ strains covers the 3’ coding region (plus stop codon and 8 bp of the 3’ UTR region), and the *nurf-1(n4295)* deletion spans 1078 bp of the 3’ coding region (Large et al. 2016). Importantly, both deletions impact the same 13 (out of a total of 16) *nurf-1* isoforms.

**Figure 8.**
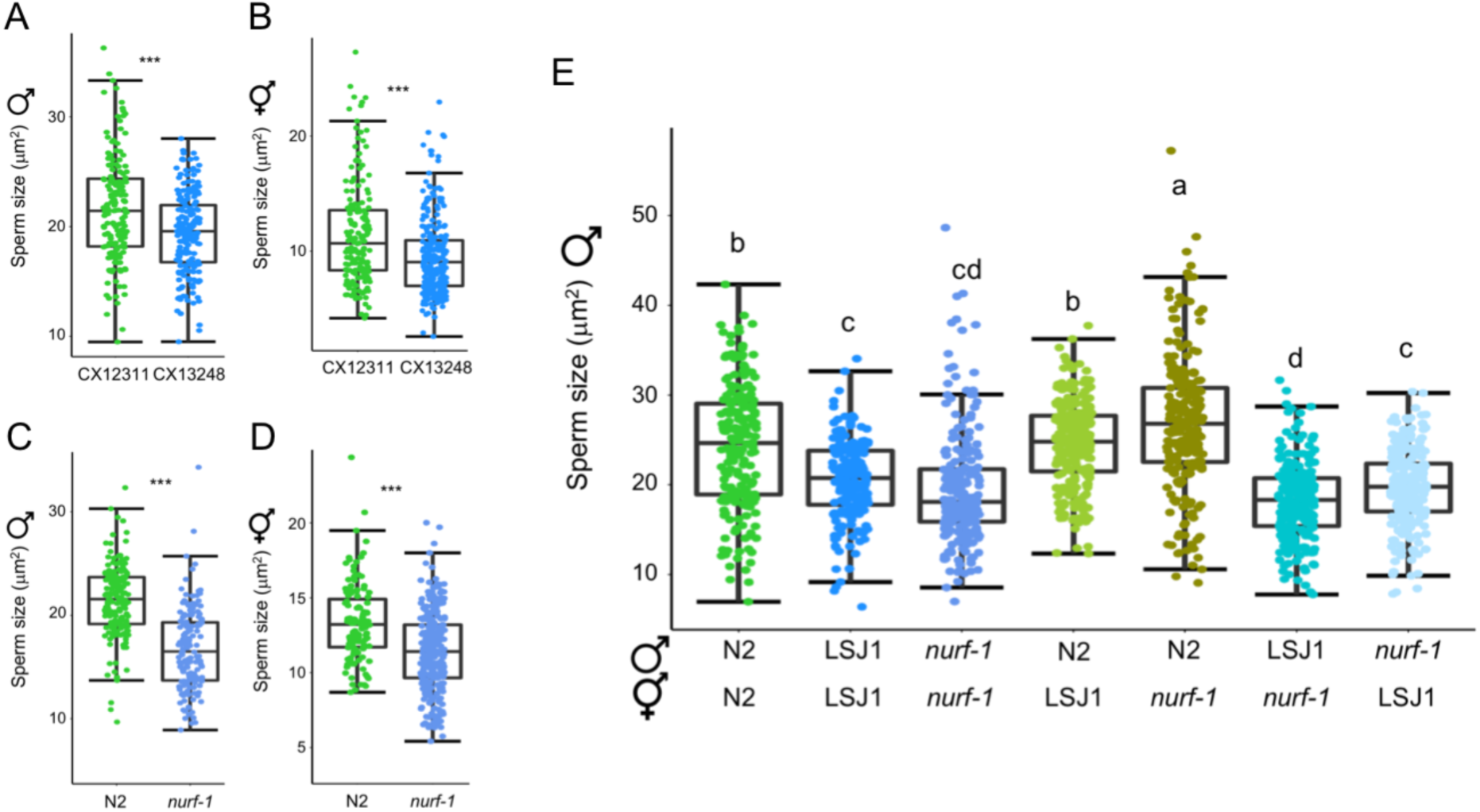
Reduced male sperm size of LSJ strains is caused by variation in *nurf-1*. (A, B) A near-isogenic line (CX13248) with the LS2 genomic region containing the *nurf-1* deletion exhibits reduced sperm size relative to the N2-like parent CX12311 in (A) males (ANOVA, F_1,336_=23.11, P<0.0001) and hermaphrodites (ANOVA, F_1,420_=30.35, P<0.0001). (C, D) Sperm size of the deletion mutant *nurf-1(n4295)* is reduced in (C) males (ANOVA, F_1,322_=184.30, P<0.0001) and (D) hermaphrodites (ANOVA, F_1,383_=61.64, P<0.0001). (E) Quantitative complementation tests using the strains N2, LSJ1, and *nurf-1(n4295)* (ANOVA, effect *strain*: F_6,1502_=97.09, P<0.0001). Values labelled with different letters indicate significant differences (Tukey’s HSD, *P* < 0.05).

To test whether *nurf-1* is the causal gene underlying reduced sperm size in the LSJ lineage, we performed a quantitative complementation test (QCT) (Long et al. 1996) taking advantage of the two *nurf-1* deletion alleles present in LSJ1 and *nurf-1(n4295)* (Figure 8E). The recessive phenotype caused by either *nurf-1* deletion was confirmed by the large, N2-like sperm size in F1 males derived from crosses between N2 and LSJ1 and between N2 and *nurf-1(n4295)* (Figure 8E). F1 males derived from bi-directional crosses between LSJ1 and *nurf-1(n4295)*, however, exhibited small sperm size, comparable to parental strains (Figure 4B). We conclude that variation in the gene *nurf-1* underlies the evolution of reduced male sperm size in LSJ strains.

Collectively, our experiments suggest that the 60 bp deletion in *nurf-1* is the causal variant responsible for the decreased sperm size in males and hermaphrodites in the LSJ lineage. First, quantitative complementation indicates *nurf-1* to be the causal gene. Second, the LSJ1/LSJ2 strains are outliers with regards to sperm size, suggesting that a mutation occurred in this lineage, like the 60bp deletion, to reduce sperm size. Finally, the *n4295* allele, which phenocopies the LSJ strains, is genetically similar to the 60 bp deletion, affecting the C-terminus of the protein. However, the intron SNV in *nurf-1*, derived in the N2 lineage, cannot be completely ruled out to explain observed sperm size differences.

### Mutants of different NURF complex components exhibit reduced sperm and spermatocyte size

Observed sperm size reduction caused by the two independent *nurf-1* deletions implies a potential role of the NURF chromatin remodelling complex in *C. elegans* sperm size determination. We therefore tested whether mutants of other complex members, including PYP-1, RBA-1 and the ATPase component ISW-1 (Figure 9A) (Tsukiyama et al. 1995; Andersen et al. 2006) show altered sperm size. Previously isolated deletion mutants of *pyp-1* and *isw-1* (Andersen et al. 2006) exhibited strongly reduced male sperm size, with *isw-1* mutants showing the most extreme sperm size reduction of all tested NURF complex mutants (Figure 9B). Using the *isw-1(n3297)* allele (Andersen et al. 2006), we further confirm that hermaphrodite sperm size was significantly reduced compared to the N2 wild-type (Figure S2).

**Figure 9.**
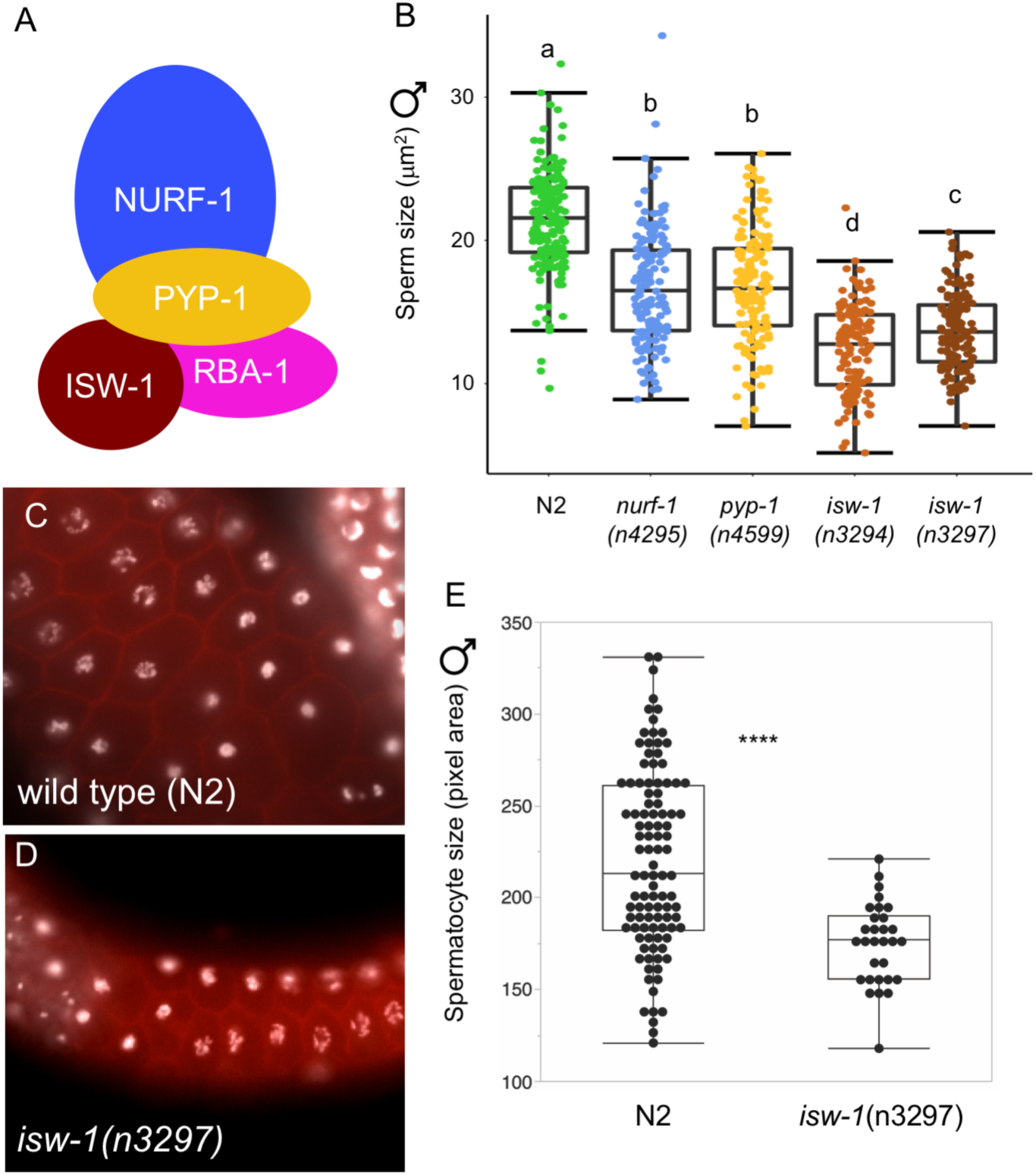
Mutants of the NURF complex exhibit reduced male sperm size and primary spermatocytes. (A) Illustration of NURF complex components. (B) Mutation of different genes in the NURF complex reduce male sperm size (ANOVA, effect *strain:* F_4,728_=192.23, P<0.0001). (C,D) Microscopy images of primary spermatocytes in (C) wild type (N2) and (D) *isw-1(n3297)* (DAPI: white, Phalloidin: red). D) Primary spermatocytes (area measurements) of the mutant *isw-1(n3297)* are significantly smaller than in the wild type (N2) (ANOVA, effect *strain*: F_1,119_=83.47, P<0.0001).

Overall germline structure and organisation of strains with small sperm size (including LSJ1 and LSJ2) appeared intact, except for a fraction of *isw-1* mutant individuals that displayed severe errors, such as displacement of spermatids into the distal region. Given that *Caenorhabditis* sperm size differences are developmentally established at the primary spermatocyte stage (Vielle et al. 2016), we measured primary male spermatocyte size in *isw-1(n3297)* animals with intact germline structure: primary spermatocyte size was on average significantly smaller than in the wild-type N2 strain (Figures 9C to 9E). Size variation of *C. elegans* sperm size observed in NURF complex mutants was thus introduced prior to, or at, the primary spermatocyte stage, as observed for sperm size variation occurring within and between different *Caenorhabditis* species (Vielle et al. 2016). Taken together, these data suggest that the NURF chromatin remodelling complex likely acts, directly or indirectly, in *C. elegans* sperm size determination. Furthermore, the role of *nurf-1* in sperm size determination seems to be evolutionarily conserved because *nurf-1* RNAi also reduced sperm size of a gonochoristic *Caenorhabditis* species, *C. plicata*, exhibiting substantially larger sperm (Vielle et al. 2016) (Figure S3).

### Male sperm size is reduced independently of body size in NURF-complex mutants

An earlier study has shown that the 60 bp deletion in *nurf-1* is responsible for reduced hermaphrodite body length in LSJ2, and similarly, the deletion allele *nurf-1(n4295)* was shown to exhibit a significantly reduced hermaphrodite body length relative to N2 (Large et al. 2016). We therefore hypothesized that perturbing activity of the NURF chromatin complex may cause systemic size reduction of diverse tissues and organs, including spermatids. Inconsistent with this hypothesis, we found male body size of *nurf-1(n4295)* with small sperm size to be significantly larger, rather than smaller than in the wild-type N2 strain (Figure 10); in addition, the *isw-1(n3294)* mutant with very small sperm had the same male body size as N2 (Figure 10). Male sperm size reduction in NURF mutants thus occurs independently of body size, suggesting that reduced male sperm size of LSJ strains is not necessarily a pleiotropic consequence of body size reduction in male body size.

**Figure 10.**
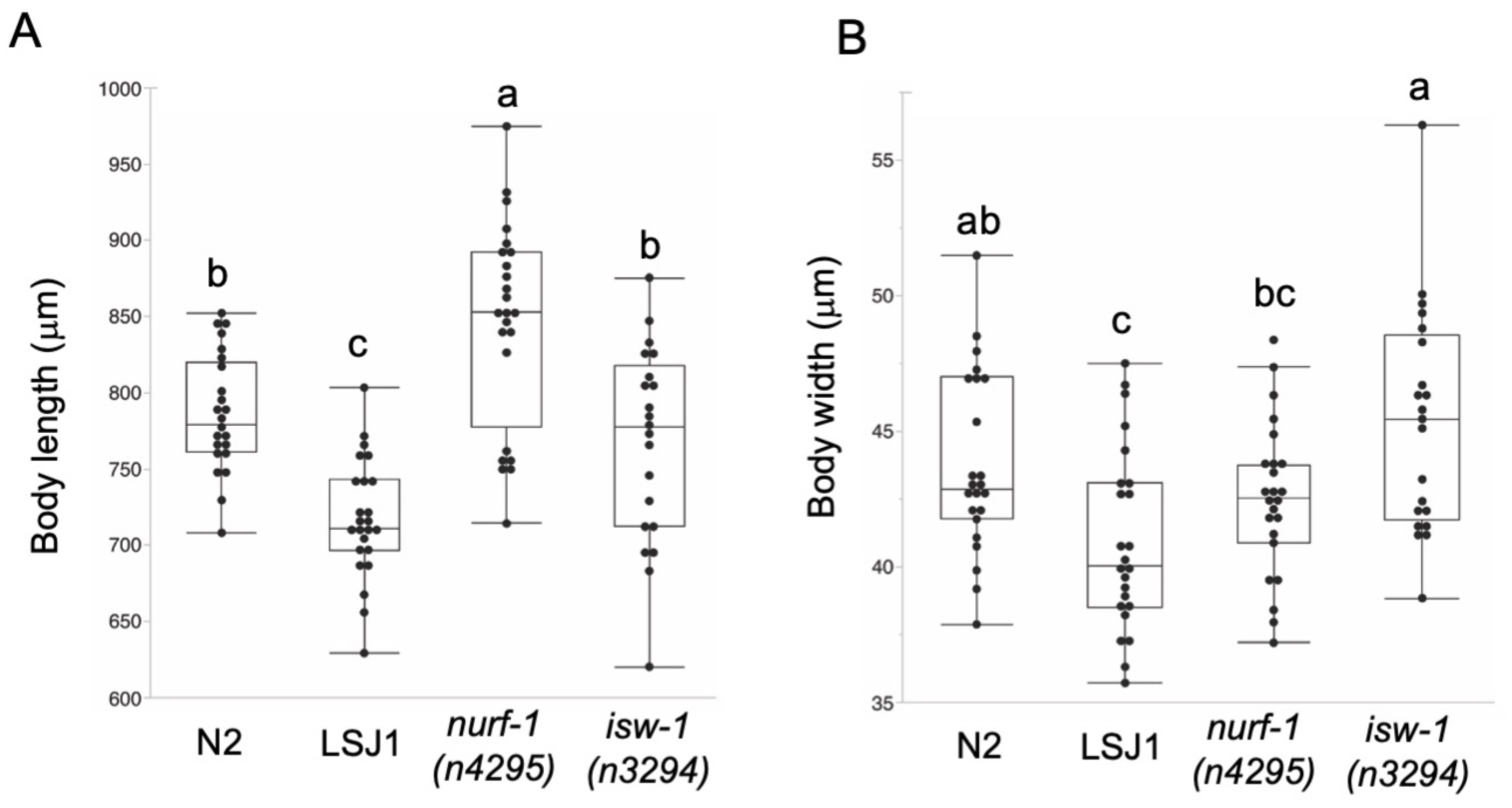
Male sperm size reduction in NURF mutants occurs independently of body size. Significant differences in body size of strains N2, LSJ1, *nurf-1(n4295)* and *isw-1(n3294).* (A) Body length (ANOVA, F_3,91_=23.36, P<0.0001). (B) Body width (ANOVA, F_3,91_=6.83, P=0.0003). Values with different letters indicate significant differences (Tukey’s HSD, *P* < 0.05).

## DISCUSSION

Our survey of intraspecific variation in *C. elegans* male sperm size uncovered significant heritable variation for this trait associated with sperm competitive ability. However, examining a subset of strains with divergent male sperm size, we did not detect a strong effect of male sperm size on competitive ability and reproductive performance. The second part of our study focused on the genetic basis underlying the evolutionary reduction in sperm size of laboratory-adapted strains, LSJ1 and LSJ2, which exhibit the smallest sperm size among all strains examined. This analysis, by means of mutants and quantitative complementation tests, shows that genetic changes in the nucleosome remodelling factor *nurf-1* underlie the evolution of small male sperm size in the LSJ lineage. These and additional experimental results suggest that the NURF chromatin remodelling complex acts in *C. elegans* sperm size regulation.

### Natural variation in *C. elegans* male sperm size

Although we observed significant heritable variation in *C. elegans* male sperm size, the vast majority of isolates show a relatively narrow average sperm size range, from 20 µm^2^ and 25 µm^2^, and the only two significant outliers were the strains JU561 and LSJ1, with reduced male sperm size (∼15 µm^2^) (Figure 1). Performing a genome-wide association study to detect potential genetic loci explaining variation in male sperm size did not yield any QTL, likely due to low statistical power and/or complex genetic trait architecture. As previously observed for some *C. elegans* isolates and other *Caenorhabditis* species (Vielle et al. 2016), male sperm size showed extreme variability within single individuals, and to a lesser extent, among populations of genetically identical individuals (Figure 1). Such within-genotype variability is often observed for sperm size traits, such as sperm length in taxa with flagellate sperm (Ward 1998; Morrow and Gage 2001b; Miller et al. 2003; Joly et al. 2004). Possibly, such high sperm size variance could reflect a means to maximize both average size and number of sperm produced, or developmentally decreasing sperm size variance may come at the cost of reduced sperm production speed (Parker and Begon 1993; Gomendio et al. 2006; Vielle et al. 2016).

### Sperm size poorly predicts male competitive ability and reproductive performance

Given the observation of heritable variation in *C. elegans* male sperm size – and the expectation that larger male sperm is more competitive – we asked whether this variation translated into differential competitive ability and reproductive success. In contrast to a previous study (Murray et al. 2011), we did not detect any significant correlations between male sperm size and male fertility or competitive ability (Figures 3 and 4). This unexpected outcome may be simply due to chance, given the small number of strains examined (n=8); nevertheless, this result shows that male sperm size is unlikely to be the only key determinant of male reproductive success, so that other traits than sperm size may be important for male fertilization success or competitive ability. Consistent with this interpretation, the number of sperm transferred per mating (ejaculate size), which also differed significantly among strains, was strongly correlated with the number of offspring sired (Figure 4F). Although we did not detect a potential trade-off between male sperm size and ejaculate size, an earlier study (Murray et al. 2011) did report that the rate of sperm production was reduced in *C. elegans* strains with larger male sperm. Trade-offs between male sperm size and sperm number are thus likely to shape and limit the extent of sperm size evolution. Together with earlier studies (Hansen et al. 2015; Vielle et al. 2016), our study suggests that size is not necessarily the central sperm trait determining male competitive ability and fertilization success. Moreover, many other traits that we did not measure here, including male mating behaviour or hermaphrodite receptivity, will contribute to overall male competitive ability and reproductive performance – as such traits may also vary across *C. elegans* strains, they could mask the relative importance of sperm size variation in determining male reproductive success.

In addition, the observed absence of a tight relationship between male sperm size and competitive ability might also represent a signature of reduced male conflict and sperm competition, in line with reports of low rates of outcrossing in *C. elegans* wild populations (Barrière and Félix 2005, 2007; Félix and Braendle 2010) and reduced maintenance of (male) mating function (Chasnov 2013; Chasnov and Chow 2002; Thomas et al. 2012; Yin et al. 2018; Teotonio et al. 2006). Nevertheless, *C. elegans* male sperm seems to maintain the evolutionary potential to respond to changes in the extent of male mating competition: several experimental evolution studies have shown that male sperm size may rapidly evolve towards increased size in response to increased male-male competition (LaMunyon and Ward 2002; Palopoli et al. 2015; Poullet et al. 2016).

### *C. elegans* sexual sperm size dimorphism

We also aimed to assess to what extent *C. elegans* sperm size differs between sexes, given that little is known about natural genetic variation in hermaphrodite sperm size and whether male and hermaphrodite sperm size may correlate. Specifically, a significant correlation between male and hermaphrodite sperm size could be indicative of shared genetic regulation of sperm size across sexes. Our analyses of intraspecific variation in *C. elegans* hermaphrodite sperm size confirmed previous reports (LaMunyon and Ward 1995, 1998; Ward and Carrel 1979; Vielle et al. 2016) that for any given strain, hermaphrodite sperm is always significantly smaller than male sperm (Figure 5A). As for male sperm size, we detected substantial levels of heritable variation in hermaphrodite sperm size. Although the two strains with smallest male sperm (LSJ1 and JU561) also had the smallest hermaphrodite sperm, male and hermaphrodite sperm size did not correlate across examined strains sperm (Figure 5). The *C. elegans* sperm size dimorphism is thought to reflect differential selection on conflicting sex-specific size optima: larger male sperm to increase sperm competitive ability versus smaller hermaphrodite sperm to accelerate sperm production to allow for a rapid switch to oogenesis, which is critical for early maturity and reproduction (Hodgkin and Barnes 1991; Cutter 2004); *C. elegans* hermaphrodites are protandrous with initial production of sperm, stored in the spermathecae, before switching to oocyte production. Therefore, the sequential spermatogenesis-oogenesis switch in *C. elegans* causes a trade-off between maximal sperm number (*i.e.* potential number of self-progeny) and minimal age at maturity (*i.e.* generation time). Consequently, evolution of small hermaphrodite sperm size results from selection for rapid self-sperm production, consistent with the fact that hermaphrodite sperm is always drastically smaller than male sperm (Figure 5A). In the same fashion, evolution of increased hermaphrodite sperm production may lead to smaller self-sperm. However, we did not find evidence for a trade-off between hermaphrodite sperm size across a set of *C. elegans* strains differing in sperm size (Figure 5C). Importantly, the evolution of small hermaphrodite sperm in androdioecious species seems not only to result from selection for improved self-fertilization but also from direct developmental effects, when spermatogenesis takes place in a female soma, that reduce hermaphrodite sperm size (Baldi et al. 2011). Disentangling these different evolutionary and developmental mechanisms in shaping *C. elegans* sperm size dimorphism therefore remains a major challenge.

### Genetic variation in *nurf-1* explains the evolution of small sperm during laboratory adaptation

Surprisingly, we uncovered extreme divergence in male sperm size between two very closely related *C. elegans* laboratory lineages, N2 versus LSJ1/LSJ2. Given their recent evolutionary divergence, and thus close genetic similarity (McGrath et al. 2011), we successfully applied a candidate approach to demonstrate that variation in the gene *nurf-1*, encoding a subunit of the NURF chromatin remodelling complex, explains reduced LSJ sperm size. The evolution of *nurf-1* function in the LSJ lineage underlies laboratory adaptation to the axenic liquid medium (Large et al. 2017). Specifically, the key causal molecular variant underlying improved LSJ hermaphrodite reproduction in this environment (relative to N2) is the 60 bp *nurf-1* deletion (Large et al. 2016), which we found to be also responsible for reduced sperm size. This deletion has been shown to have highly pleiotropic effects on life history traits, including reproduction, growth rate, life span and dauer formation (Large et al. 2016). Our result suggests that these life history changes may be, at least partly, mediated by changes in *nurf-1* function in the germline, affecting sperm size, likely also in hermaphrodites. Furthermore, evolution of reduced hermaphrodite sperm size could thus be directly linked to the observed evolutionary increase in hermaphrodite sperm number (Figure 5C), although this remains to be experimentally verified; we did not formally show that the reduced hermaphrodite sperm size is due to the same genetic variant(s) in *nurf-1* as in males. Given the previous demonstration that *nurf-1* variants specific to the LSJ lineage confer improved fitness of hermaphrodites, the evolution of reduced male sperm size in the LSJ lineage likely reflects a pleiotropic consequence stemming from selection on hermaphrodite function. Moreover, because *C. elegans* males are generally incapable of mating in liquid culture, it appears likely that laboratory evolution of the LSJ lineage likely occurred in the complete absence of outcrossing. This result illustrates how selection for improved *C. elegans* hermaphrodite function (Large et al. 2016) can cause the deterioration of male-specific fitness trait (sperm size) through a specific sexually antagonistic genetic variant. Evolutionary variation at the *nurf-1* gene can thus be considered a potential example of intralocus sexual conflict (Bonduriansky and Chenoweth 2009). Consistent with this interpretation, we observed that LSJ1 showed significantly reduced male-male competitive ability relative to N2 (Figures 4C and 4E). On the other hand, despite the presumed absence of outcrossing over many hundreds of generations in liquid culture, we also found that LSJ1 male sperm function and male mating ability remained largely preserved (Figures 3 and 4). These observations exemplify how essential male functions in *C. elegans* can be maintained despite very rare outcrossing, *i.e.* strongly relaxed selection, due to pleiotropic effects, in particular, selection on hermaphrodite sperm function as sperm genes are shared by both sexes.

### A role for the NURF chromatin remodelling complex in *C. elegans* sperm size determination

The observation that evolutionary reduction of *C. elegans* male sperm size is caused by variation in *nurf-1* suggests that this gene acts in *C. elegans* sperm size determination. Our analysis of multiple mutants in NURF-complex genes (*nurf-1, isw-1, pyp-1*), all of which displayed strongly reduced sperm size, not only in males but also in hermaphrodites, confirms this idea. As a subunit of the NURF chromatin remodelling complex, NURF-1 is a BPTF orthologue, involved in histone modification on nucleosomes and remodelling of nucleosomes through recruitment of ISWI, to modulate transcription (Badenhorst et al. 2002; Andersen et al. 2006; Ruthenburg et al. 2011; Wysocka et al. 2006). Members of the *C. elegans* NURF chromatin remodelling complex are expressed in diverse tissues and organs, including the developing germline (Craig et al. 2013; Feng et al. 2012; Reece-Hoyes et al. 2007). Consistent with these expression patterns, *nurf-1* mutations affect diverse phenotypes in *C. elegans*, ranging from vulval development (Andersen et al. 2006) to multiple life history traits (Large et al. 2016). Therefore, whether the NURF chromatin remodelling complex specifically acts in the *C. elegans* germline to regulate expression of target genes modulating sperm size remains to be tested. Nevertheless, given the combined experimental evidence that NURF-1 activity affects (a) sperm size (this study), (b) *C. briggsae* spermatogenesis (Chen et al. 2014), and (c) reproductive timing and number of self-progeny (Large et al. 2016) clearly points to a specific role of the NURF chromatin remodelling in *C. elegans* sperm phenotypes.

## ACKNOWLEDGEMENTS

We thank Asher Cutter, Marie-Anne Félix and Henrique Teotonio for discussion and comments on previous versions of the manuscript. Strains and materials were provided by the groups of Julie Ahringer, Cori Bargmann, Marie-Anne Félix, the *C. elegans* Natural Diversity Resource (CeNDR) (Cook et al. 2017), and the *Caenorhabditis* Genetics Center (CGC), which is funded by NIH Office of Research Infrastructure Programs (P40 OD010440). CB, CG and AV acknowledge financial support by the Centre National de la Recherche Scientifique (CNRS). NS was supported by an Erasmus mobility fellowship provided by the European Commission. SZ was supported by the National Institutes of Health Cell and Molecular Basis of Disease training grant (T32GM008061) and a Bernard and Martha Rappaport Fellowship. PTM was supported by NIH grant R01GM114170. ECA was supported by an American Cancer Society Research Scholar Grant (127313-RSG-15-135-01-DD).

## SUPPORTING INFORMATION

Evolution of sperm competition: Natural variation and genetic determinants of *Caenorhabditis elegans* sperm size by Gimond *et al.*

### Supporting Files

**File S1. Raw data in Excel format. Numbers below refer to different worksheets in Excel file**

1. Male sperm size of 97 *C. elegans* strains (measurements in microns) (Figure 1)
2. Strain variation in male mating ability and fertility (Figures 3A and 3C)
3. Strain variation in ejaculate size (sperm number) (Figure 3E)
4. Strain variation in male fertilization success when competing with hermaphrodite self-sperm (Figure 4A)
5. Strain variation in male-male competitive ability (Figures 4C and 4E)
6. Strain variation in hermaphrodite sperm size (Figure 5A)
7. Strain variation in hermaphrodite sperm production (Figure 5C)
8. Strain variation in body size (Figure 6)
9. Male and hermaphrodite sperm size of strains N2, LSJ1, LSJ2 and CC1 (Figure 7)
10. Male and hermaphrodite sperm size of strains CX12311, CX13248, N2, and *nurf-1(n4295)* (Figure 8A-D)
11. Male sperm size measurements resulting from quantitative complementation tests using strains N2, LSJ1 and *nurf-1(n4295)* (Figure 8E)
12. Male sperm size of NURF complex mutants (Figure 9B)
13. Hermaphrodite sperm size of *isw-1(n3297)* versus N2 (Figure S2)
14. Male primary spermatocyte size of *isw-1(n3297)* versus N2 (Figure 9E)
15. Variation in body size of N2, LSJ1, *nurf-1(n4295)* and *isw-1(n3294)* (Figure 10)
16. Effect of *nurf-1* RNAi on male sperm size of *C. elegans* (Figure S1)
17. Effect of *nurf-1* RNAi on male sperm size of *C. plicata* (Figure S3)

### Supporting Figures

**Figure S1.**
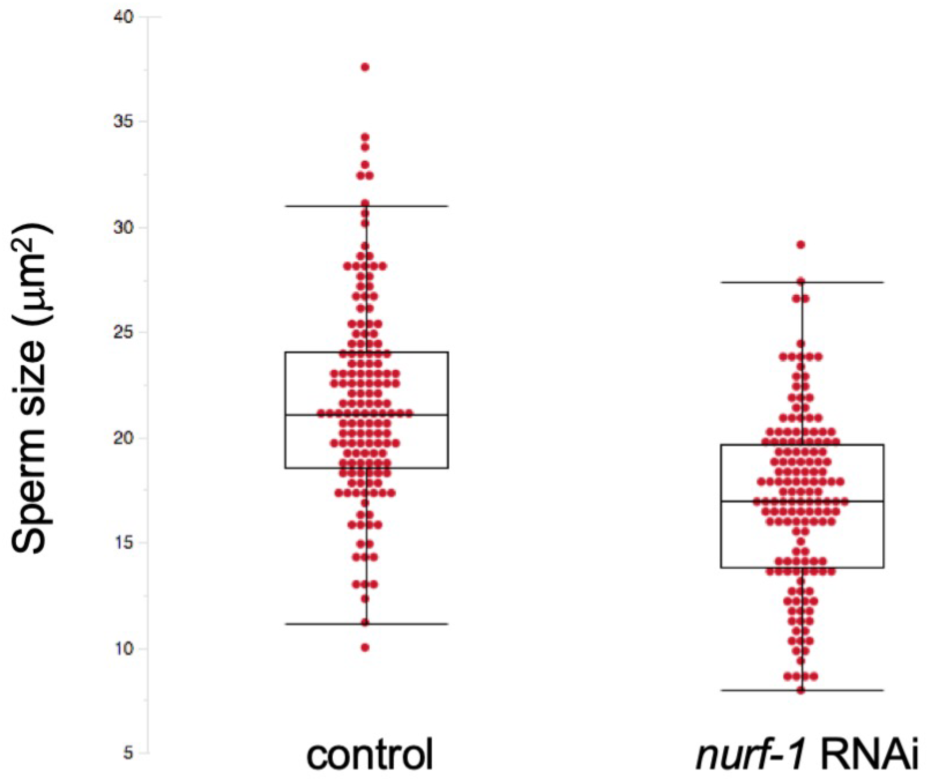
Effect of *nurf-1* RNAi on male sperm size of *C. elegans*. *nurf-1* RNAi significantly reduces male sperm size of the reference strain N2 (ANOVA, effect *treatment*: F_1, 280_ =90.91, P<0.0001; effect *individual(treatment)*: F_18, 280_ =3.07, P<0.0001). Fifteen spermatids from each of 10 individuals were measured per treatment (N=150).

**Figure S2.**
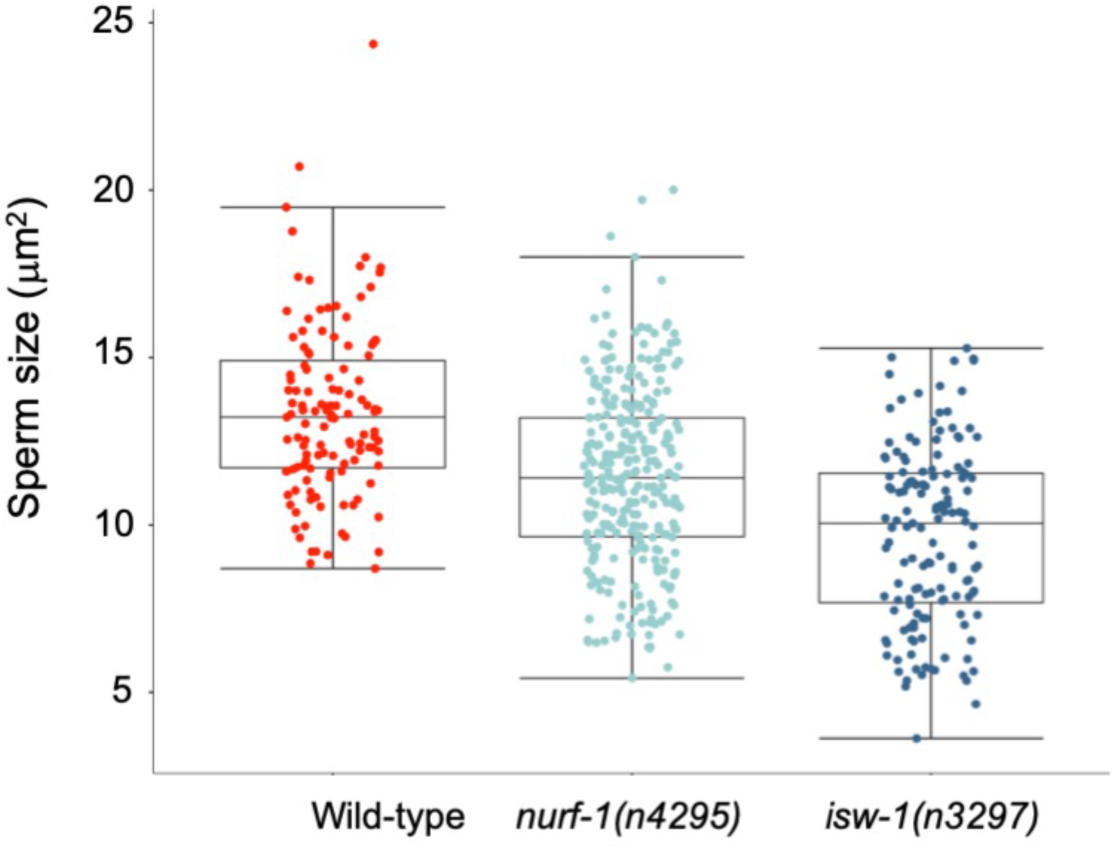
Hermaphrodite sperm size of *isw-1(n3297)* compared to N2 wild-type and *nurf-1(n4295).* Hermaphrodite sperm size of *isw-1(n3297)* is significantly reduced compared to the N2 wild-type (ANOVA, effect *strain*: F_1, 280_ =90.91, P<0.0001; effect *individual(strain)*: F_18, 280_ =3.07, P<0.0001).

**Figure S3.**
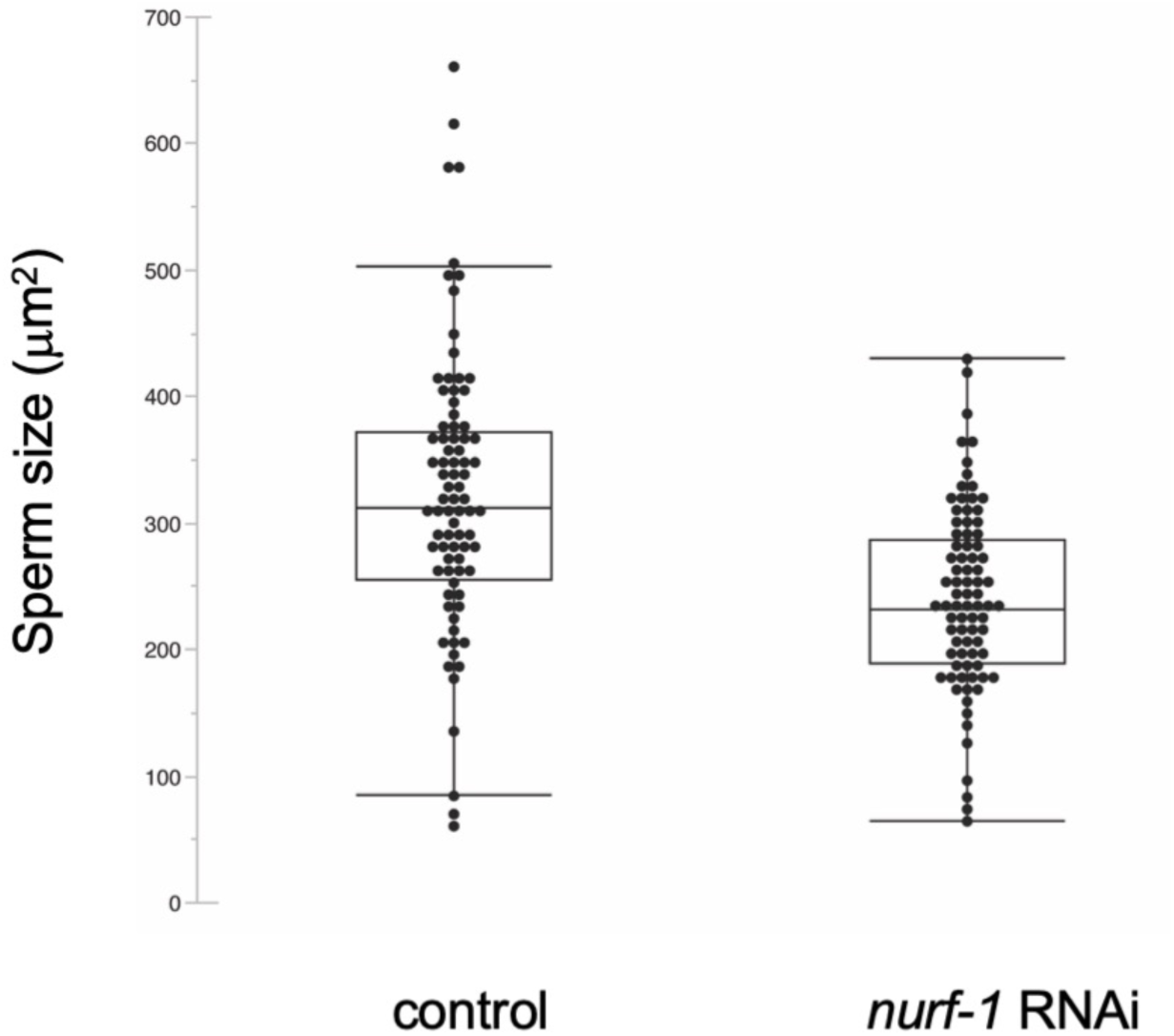
Effect of *nurf-1* RNAi on male sperm size of the gonochoristic species, *C. plicata.* *nurf-1 (C. elegans)* RNAi significantly reduces male sperm size of the strain SB355 (ANOVA, effect *treatment*: F_1, 163_ =27.56, P<0.0001; effect *individual(treatment)*: F_10,163_ =8.10, P<0.0001). 8-15 spermatids from each of 7 individuals were measured per treatment.

### Supporting Tables

**Table S1.** List of *Caenorhabditis* strains used in this study. For additional strain information, see CeNDR: https://www.elegansvariation.org)

| STRAIN | ORIGIN | DATE OF ISOLATION |
| --- | --- | --- |
| AB1 | Adelaide, Australia | 1983 |
| AB4 | Adelaide, Australia | 1983 |
| CB4851 | Bergerac, France | pre-1949 |
| CB4852 | Unknown | pre-1966 |
| CB4853 | Altadena, USA | 1974/05 |
| CB4854 | Altadena, USA | 1974/05 |
| CB4856 | Oahu, Hawaii | 1972/08 |
| CB4857 | Claremont, USA | 1972/11 |
| CB4858 | Pasadena, USA | 1973 |
| CB4932 | Taunton, United Kingdom | pre-1991 |
| CX11262 | Los Angeles, USA | 2003/09 |
| CX11264 | Los Angeles, USA | 2003/09 |
| CX11271 | Pasadena, USA | 2003/09 |
| CX11276 | Los Angeles, USA | 2003/09 |
| CX11285 | Los Angeles, USA | 2003/09 |
| CX11292 | Los Angeles, USA | 2004/02 |
| CX11307 | Los Angeles, USA | 2003/09 |
| CX11314 | Los Angeles, USA | 2003/09 |
| CX11315 | Los Angeles, USA | 2003/09 |
| DL200 | Addis Ababa, Ethiopia | 2007/12 |
| DL226 | Corvallis, USA | 2007 |
| DL238 | Manuka, Hawaii | 2008/07/15 |
| ED3005 | Edinburgh, United Kingdom | 2004/10/25 |
| ED3011 | Edinburgh, United Kingdom | 2004/11/26 |
| ED3012 | Edinburgh, United Kingdom | 2004/11/26 |
| ED3017 | Edinburgh, United Kingdom | 2004/12/03 |
| ED3040 | Johannesburg, South Africa | 2006/03 |
| ED3046 | Ceres, South Africa | 2006/03 |
| ED3048 | Ceres, South Africa | 2006/03 |
| ED3049 | Ceres, South Africa | 2006/03 |
| ED3052 | Ceres, South Africa | 2006/03 |
| ED3073 | Limuru, Kenya | 2006/03 |
| ED3077 | Nairobi, Kenya | 2006/03 |
| EG4347 | Eugene, USA | 2006/10 |
| EG4349 | Salt Lake City, USA | 2006/10 |
| EG4724 | Amares, Portugal | 2007/03 |
| EG4725 | Amares, Portugal | 2007/03 |
| EG4946 | Salt Lake City, USA | 2007/09/27 |
| JT11398 | Lake Forest Park, USA | 2003/12 |
| JU1088 | Kakegawa, Japan | 2007/03/14 |
| JU1172 | Concepcion, Chile | 2007/04 |
| JU1200 | Dundonald, United Kingdom | 2007/08/01 |
| JU1212 | Primel Trégastel, France | 2007/09/24 |
| JU1213 | Primel Trégastel, France | 2007/09/24 |
| JU1242 | Santeuil, France | 2007/10/14 |
| JU1246 | Santeuil, France | 2007/10/14 |
| JU1395 | Saut-aux-Loups, France | 2008/03/01 |
| JU1400 | Sevilla, Spain | 2008/03 |
| JU1409 | Carmona, Spain | 2008/03/31 |
| JU1440 | Barcelona, Spain | 2008/06/09 |
| JU1491 | Le Blanc, France | 2008/08/17 |
| JU1530 | Orsay, France | 2008/09/09 |
| JU1568 | Ivry-sur-Seine, France | 2008/10/05 |
| JU1580 | Orsay, France | 2008/10/06 |
| JU1581 | Orsay, France | 2008/10/23 |
| JU1586 | Le Blanc, France | 2008/11/03 |
| JU1652 | Montevideo, Uruguay | 2009 |
| JU1896 | Athens, Greece | 2010/01/02 |
| JU258 | Ribeiro Frio, Madeira | 2001/10 |
| JU310 | Le Blanc, France | 2002/08/25 |
| JU311 | Merlet, France | 2002/09/08 |
| JU323 | Merlet, France | 2002/09/08 |
| JU346 | Merlet, France | 2002/09/08 |
| JU360 | Franconville, France | 2002/09/02 |
| JU363 | Franconville, France | 2002/09/16 |
| JU367 | Franconville, France | 2002/09/16 |
| JU393 | Hermanville, France | 2002/09 |
| JU394 | Hermanville, France | 2002/09 |
| JU397 | Hermanville, France | 2002/09 |
| JU406 | Hermanville, France | 2002/12/30 |
| JU440 | Beauchêne, France | 2003/09/12 |
| JU561 | Sainte-Barbe, France | 2004/10/03 |
| JU642 | Le Perreux-sur-Marne, France | 2004/12/14 |
| JU751 | Le Perreux-sur-Marne, France | 2005/06/08 |
| JU774 | Carcavelos, Portugal | 2005/07/10 |
| JU775 | Lisbon, Portugal | 2005/07/10 |
| JU778 | Lisbon, Portugal | 2005/07/10 |
| JU782 | Lisbon, Portugal | 2005/07/10 |
| JU792 | Fréchendets, France | 2005/08/31 |
| JU830 | Tübingen, Germany | 2005/09/28 |
| JU847 | Obernai, France | 2005/10/03 |
| KR314 | Vancouver, Canada | 1984/05 |
| LKC34 | Unknown city, Madagascar | 2005/06/17 |
| LSJ1 | Bristol, United Kingdom | 1951 |
| N2 | Bristol, United Kingdom | 1951 |
| MY1 | Lingen, Germany | 2002/07 |
| MY10 | Roxel, Germany | 2002/07 |
| MY16 | Mecklenbeck, Germany | 2002/07 |
| MY18 | Roxel, Germany | 2002/07 |
| MY23 | Roxel, Germany | 2002/07 |
| PB303 | NA | 1998/11/14 |
| PB306 | NA | 1998/11/28 |
| PS2025 | Altadena, USA | Unknown |
| PX179 | Eugene, USA | 2001/10/02 |
| QX1211 | San Francisco, USA | 2007/11/26 |
| QX1233 | Berkeley, USA | 2007/11/24 |
| RC301 | Freiburg, Germany | 1983 |
| WN2002 | Wageningen, Netherlands | 2007/11/20 |

**Table S2.** Male sperm size variation (means of cross-sectional area, microns) and CV in 97 *C. elegans* strains.

| Strain | Sperm Size (mean) | SEM | CV (within-individual) | CV (between-individual) | N individuals | N sperm |
| --- | --- | --- | --- | --- | --- | --- |
| AB1 | 23.94 | 1.68 | 16.49 | 5.22 | 7 | 140 |
| AB4 | 22.13 | 0.83 | 18.75 | 6.46 | 7 | 140 |
| CB4852 | 21.44 | 1.91 | 17.55 | 5.68 | 7 | 140 |
| CB4853 | 21.98 | 2.36 | 22.13 | 4.66 | 7 | 140 |
| CB4854 | 25.07 | 1.73 | 21.09 | 4.05 | 7 | 140 |
| CB4856 | 22.83 | 0.70 | 16.42 | 5.60 | 7 | 140 |
| CB4857 | 22.89 | 2.05 | 20.67 | 4.35 | 7 | 140 |
| CB4858 | 22.08 | 1.53 | 17.63 | 7.01 | 7 | 140 |
| CB4932 | 21.06 | 0.88 | 15.65 | 7.69 | 7 | 140 |
| CX11262 | 21.81 | 1.65 | 16.93 | 8.79 | 7 | 140 |
| CX11264 | 20.85 | 1.29 | 16.33 | 4.91 | 7 | 140 |
| CX11271 | 25.03 | 1.55 | 18.86 | 7.04 | 7 | 140 |
| CX11276 | 21.56 | 1.24 | 18.82 | 6.23 | 7 | 140 |
| CX11285 | 18.97 | 1.60 | 17.16 | 6.81 | 7 | 140 |
| CX11292 | 21.40 | 1.51 | 19.59 | 7.48 | 7 | 140 |
| CX11307 | 23.95 | 1.82 | 17.03 | 5.90 | 7 | 140 |
| CX11314 | 20.76 | 1.83 | 18.55 | 6.06 | 7 | 140 |
| CX11315 | 20.82 | 1.36 | 15.20 | 7.69 | 7 | 140 |
| DL200 | 20.62 | 0.62 | 16.97 | 3.31 | 7 | 140 |
| DL226 | 24.36 | 2.09 | 18.99 | 5.82 | 7 | 140 |
| DL238 | 21.29 | 1.52 | 19.70 | 8.92 | 7 | 140 |
| ED3005 | 24.36 | 1.47 | 24.36 | 5.44 | 7 | 140 |
| ED3011 | 21.17 | 1.39 | 25.29 | 13.02 | 7 | 140 |
| ED3012 | 22.72 | 1.05 | 16.88 | 6.27 | 7 | 140 |
| ED3017 | 23.23 | 1.70 | 20.66 | 4.92 | 7 | 140 |
| ED3040 | 22.45 | 1.14 | 20.66 | 9.96 | 7 | 140 |
| ED3046 | 24.32 | 1.22 | 18.63 | 4.75 | 7 | 140 |
| ED3048 | 20.73 | 2.29 | 19.68 | 6.12 | 7 | 140 |
| ED3049 | 21.68 | 0.72 | 13.74 | 4.87 | 7 | 140 |
| ED3052 | 21.97 | 1.91 | 16.82 | 7.09 | 7 | 140 |
| ED3073 | 22.98 | 1.45 | 16.42 | 5.25 | 7 | 140 |
| ED3077 | 22.63 | 0.79 | 20.15 | 6.86 | 7 | 140 |
| EG4347 | 24.05 | 1.28 | 19.25 | 7.10 | 7 | 140 |
| EG4349 | 21.49 | 2.00 | 18.71 | 5.43 | 7 | 140 |
| EG4724 | 20.96 | 2.24 | 14.97 | 6.88 | 7 | 140 |
| EG4725 | 21.08 | 1.24 | 18.07 | 8.55 | 7 | 140 |
| EG4946 | 25.55 | 1.68 | 19.75 | 6.28 | 7 | 140 |
| JT11398 | 22.17 | 1.80 | 15.50 | 5.90 | 7 | 140 |
| JU258 | 20.79 | 1.46 | 15.15 | 5.98 | 7 | 140 |
| JU310 | 24.36 | 1.23 | 19.54 | 5.79 | 7 | 140 |
| JU311 | 24.06 | 2.07 | 20.87 | 13.30 | 7 | 140 |
| JU323 | 23.80 | 2.55 | 22.45 | 6.77 | 7 | 140 |
| JU346 | 23.35 | 1.66 | 14.92 | 5.96 | 7 | 140 |
| JU360 | 23.91 | 1.34 | 17.15 | 7.36 | 7 | 140 |
| JU363 | 24.90 | 1.38 | 19.22 | 5.31 | 7 | 140 |
| JU367 | 23.37 | 0.94 | 14.16 | 5.77 | 7 | 140 |
| JU393 | 26.70 | 0.92 | 18.15 | 10.12 | 7 | 140 |
| JU394 | 24.16 | 1.39 | 16.32 | 4.12 | 7 | 140 |
| JU397 | 23.12 | 0.95 | 12.10 | 6.74 | 6 | 120 |
| JU406 | 21.36 | 1.96 | 17.72 | 7.32 | 7 | 140 |
| JU440 | 23.22 | 1.01 | 17.77 | 6.27 | 7 | 140 |
| JU561 | 15.92 | 2.06 | 19.17 | 12.52 | 7 | 140 |
| JU642 | 21.21 | 1.55 | 13.97 | 11.23 | 7 | 140 |
| JU751 | 21.94 | 2.41 | 18.75 | 9.02 | 7 | 140 |
| JU774 | 22.10 | 1.68 | 21.79 | 10.10 | 7 | 140 |
| JU775 | 22.48 | 1.89 | 21.71 | 10.49 | 7 | 140 |
| JU778 | 20.37 | 1.46 | 22.84 | 8.98 | 7 | 140 |
| JU782 | 26.29 | 2.68 | 23.26 | 3.23 | 7 | 140 |
| JU792 | 20.59 | 1.40 | 14.52 | 7.78 | 7 | 140 |
| JU830 | 24.16 | 1.06 | 15.54 | 3.07 | 7 | 140 |
| JU847 | 22.06 | 1.73 | 17.73 | 5.16 | 7 | 140 |
| JU1088 | 23.41 | 1.19 | 14.26 | 8.82 | 7 | 140 |
| JU1172 | 22.57 | 2.48 | 20.17 | 5.61 | 7 | 140 |
| JU1200 | 22.00 | 1.37 | 18.35 | 3.85 | 7 | 140 |
| JU1212 | 21.61 | 1.51 | 18.11 | 3.14 | 7 | 140 |
| JU1213 | 23.80 | 1.44 | 24.69 | 21.07 | 7 | 140 |
| JU1242 | 19.99 | 1.00 | 15.13 | 12.12 | 7 | 140 |
| JU1246 | 21.40 | 2.59 | 19.01 | 7.33 | 7 | 140 |
| JU1395 | 22.77 | 1.04 | 17.56 | 2.38 | 7 | 140 |
| JU1400 | 19.56 | 1.91 | 18.74 | 5.19 | 7 | 140 |
| JU1409 | 24.64 | 0.91 | 13.12 | 5.23 | 7 | 140 |
| JU1440 | 20.49 | 2.44 | 19.04 | 4.92 | 7 | 140 |
| JU1491 | 21.70 | 1.71 | 21.68 | 3.81 | 7 | 140 |
| JU1530 | 25.71 | 1.63 | 17.35 | 20.12 | 7 | 140 |
| JU1568 | 22.09 | 1.91 | 16.93 | 6.42 | 7 | 140 |
| JU1580 | 22.00 | 0.76 | 15.02 | 7.87 | 7 | 140 |
| JU1581 | 20.41 | 1.22 | 13.76 | 3.65 | 7 | 140 |
| JU1586 | 20.16 | 1.47 | 19.50 | 9.14 | 7 | 140 |
| JU1652 | 21.95 | 1.59 | 17.23 | 6.97 | 7 | 140 |
| JU1896 | 23.21 | 0.81 | 18.26 | 8.87 | 7 | 140 |
| KR314 | 22.11 | 3.11 | 21.29 | 14.20 | 6 | 120 |
| LKC34 | 24.68 | 1.06 | 18.99 | 5.75 | 7 | 140 |
| LSJ1 | 15.25 | 0.82 | 16.09 | 9.08 | 7 | 140 |
| MY1 | 20.01 | 1.56 | 16.72 | 4.62 | 7 | 140 |
| MY10 | 20.84 | 1.54 | 20.29 | 9.43 | 7 | 140 |
| MY16 | 22.46 | 1.79 | 16.80 | 1.66 | 7 | 140 |
| MY18 | 22.16 | 1.12 | 20.60 | 10.50 | 7 | 140 |
| MY23 | 20.09 | 1.65 | 16.80 | 7.44 | 7 | 140 |
| N2 | 19.51 | 2.17 | 20.68 | 3.70 | 7 | 140 |
| PB303 | 23.01 | 0.83 | 18.10 | 4.90 | 7 | 140 |
| PB306 | 22.15 | 1.47 | 18.60 | 5.82 | 7 | 140 |
| PS2025 | 22.40 | 1.48 | 19.73 | 7.99 | 7 | 140 |
| PX179 | 22.41 | 1.81 | 21.73 | 7.20 | 7 | 140 |
| QX1211 | 22.11 | 1.02 | 18.49 | 6.97 | 7 | 140 |
| QX1233 | 23.57 | 2.81 | 23.73 | 19.28 | 7 | 140 |
| RC301 | 21.86 | 0.95 | 16.92 | 6.65 | 7 | 140 |
| WN2002 | 22.37 | 2.06 | 20.86 | 7.31 | 7 | 140 |

**Table S3.** Mean sperm size and plugging phenotype of examined *C. elegans* strains. Plugging phenotype data from Andersen et al. (2012) and Palopoli et al. (2015).

| Strain | Mean sperm size | Plugging =1, Non-plugging=0 |
| --- | --- | --- |
| AB1 | 23.5511 | 0 |
| AB4 | 21.659 | 1 |
| CB4852 | 21.0393 | 0 |
| CB4853 | 21.356 | 1 |
| CB4854 | 24.4207 | 1 |
| CB4856 | 22.4802 | 1 |
| CB4857 | 22.3462 | 0 |
| CB4858 | 21.6617 | 1 |
| CB4932 | 20.7466 | 0 |
| CX11262 | 21.3944 | 1 |
| CX11264 | 20.537 | 1 |
| CX11271 | 24.5216 | 1 |
| CX11276 | 21.1239 | 1 |
| CX11285 | 18.6448 | 1 |
| CX11292 | 20.9053 | 1 |
| CX11307 | 23.5335 | 1 |
| CX11314 | 20.3464 | 1 |
| CX11315 | 20.5113 | 1 |
| DL200 | 20.3174 | 1 |
| DL226 | 23.8156 | 1 |
| DL238 | 20.7834 | 1 |
| ED3005 | 23.5608 | 1 |
| ED3011 | 20.2955 | 1 |
| ED3012 | 22.3289 | 0 |
| ED3017 | 22.6823 | 1 |
| ED3040 | 21.8398 | 1 |
| ED3046 | 23.8324 | 1 |
| ED3048 | 20.2543 | 1 |
| ED3049 | 21.45 | 1 |
| ED3052 | 21.5752 | 1 |
| ED3073 | 22.6221 | 1 |
| ED3077 | 22.0667 | 1 |
| EG4347 | 23.5467 | 0 |
| EG4349 | 21.0612 | 1 |
| EG4724 | 20.6426 | 1 |
| EG4725 | 20.631 | 1 |
| EG4946 | 24.9506 | 0 |
| JT11398 | 21.8455 | 0 |
| JU1088 | 23.0792 | 1 |
| JU1172 | 22.0359 | 1 |
| JU1200 | 21.58 | 0 |
| JU1212 | 21.2382 | 0 |
| JU1213 | 22.6857 | 1 |
| JU1242 | 19.6241 | 1 |
| JU1246 | 20.9496 | 1 |
| JU1395 | 22.3787 | 0 |
| JU1400 | 19.1491 | 1 |
| JU1409 | 24.3819 | 1 |
| JU1440 | 20.0642 | 1 |
| JU1491 | 21.1306 | 1 |
| JU1530 | 24.8259 | 1 |
| JU1568 | 21.6964 | 0 |
| JU1580 | 21.6949 | 1 |
| JU1581 | 20.1968 | 1 |
| JU1586 | 19.6909 | 0 |
| JU1652 | 21.5463 | 1 |
| JU1896 | 22.7062 | 0 |
| JU258 | 20.4938 | 1 |
| JU310 | 23.8629 | 0 |
| JU311 | 23.3055 | 0 |
| JU323 | 23.1108 | 1 |
| JU346 | 23.0378 | 1 |
| JU360 | 23.4768 | 1 |
| JU363 | 24.3861 | 1 |
| JU367 | 23.1158 | 0 |
| JU393 | 26.1593 | 1 |
| JU394 | 23.8254 | 0 |
| JU397 | 22.9068 | 1 |
| JU406 | 20.9359 | 0 |
| JU440 | 22.8045 | 0 |
| JU561 | 15.4953 | 1 |
| JU642 | 20.8737 | 1 |
| JU751 | 21.4432 | 1 |
| JU774 | 21.4306 | 1 |
| JU775 | 21.8151 | 1 |
| JU778 | 19.7247 | 1 |
| JU782 | 25.4637 | 1 |
| JU792 | 20.3049 | 1 |
| JU830 | 23.8397 | 1 |
| JU847 | 21.6807 | 1 |
| KR314 | 21.366 | 1 |
| LKC34 | 24.1826 | 1 |
| LSJ1 | 15.0017 | 0 |
| MY1 | 19.7027 | 1 |
| MY10 | 20.3428 | 1 |
| MY16 | 22.107 | 1 |
| MY18 | 21.557 | 1 |
| MY23 | 19.733 | 1 |
| N2 | 19.0539 | 0 |
| PB303 | 22.6011 | 1 |
| PB306 | 21.7096 | 1 |
| PS2025 | 21.8714 | 1 |
| PX179 | 21.788 | 0 |
| QX1211 | 21.6496 | 1 |
| QX1233 | 22.4217 | 1 |
| RC301 | 21.4845 | 1 |
| WN2002 | 21.7914 | 0 |

**Table S4.** Strain variation in hermaphrodite sperm size (and comparison to male sperm size) (cross-sectional area in microns) (data shown in figure 5A).

| Strain | Sex | Mean sperm size | SEM | N individuals | N spermatids |
| --- | --- | --- | --- | --- | --- |
| LSJ1 | Male | 15.25 | 0.23 | 7 | 140 |
| LSJ1 | Hermaphrodite | 10.44 | 0.20 | 11 | 143 |
| JU561 | Male | 15.91 | 0.31 | 7 | 140 |
| JU561 | Hermaphrodite | 10.34 | 0.17 | 12 | 134 |
| CX11285 | Male | 18.97 | 0.29 | 7 | 140 |
| CX11285 | Hermaphrodite | 12.04 | 0.22 | 9 | 104 |
| N2 | Male | 19.51 | 0.35 | 7 | 140 |
| N2 | Hermaphrodite | 13.37 | 0.23 | 9 | 123 |
| DL238 | Male | 21.29 | 0.39 | 7 | 140 |
| DL238 | Hermaphrodite | 9.96 | 0.19 | 8 | 140 |
| JU751 | Male | 21.94 | 0.38 | 7 | 140 |
| JU751 | Hermaphrodite | 11.65 | 0.25 | 10 | 99 |
| JU1200 | Male | 21.99 | 0.34 | 7 | 140 |
| JU1200 | Hermaphrodite | 12.16 | 0.20 | 12 | 136 |
| JU775 | Male | 22.48 | 0.46 | 7 | 140 |
| JU775 | Hermaphrodite | 9.04 | 0.24 | 7 | 87 |
| CB4856 | Male | 22.8 | 0.33 | 7 | 140 |
| CB4856 | Hermaphrodite | 13.25 | 0.21 | 13 | 149 |
| EG4946 | Male | 25.55 | 0.44 | 7 | 140 |
| EG4946 | Hermaphrodite | 11.07 | 0.25 | 8 | 125 |
| JU782 | Male | 26.29 | 0.53 | 7 | 140 |
| JU782 | Hermaphrodite | 12.94 | 0.24 | 10 | 136 |
| JU393 | Male | 26.70 | 0.45 | 7 | 140 |
| JU393 | Hermaphrodite | 13.19 | 0.32 | 7 | 152 |

**Table S5.** Strain variation in hermaphrodite sperm production (data in Figure 5C).

| **Strain** | **Mean sperm**<br>**number/spermatheca** | **SEM** | **N individuals** |
| --- | --- | --- | --- |
| CB4856 | 137.00 | 3.28 | 31 |
| CX11285 | 146.65 | 3.98 | 20 |
| EG4946 | 151.83 | 7.22 | 23 |
| JU393 | 150.15 | 5.48 | 20 |
| JU561 | 136.38 | 5.20 | 29 |
| JU782 | 152.74 | 4.00 | 27 |
| LSJ1 | 175.96 | 7.81 | 23 |
| N2 | 145.00 | 5.28 | 12 |

**Table S6.** Strain and sex differences in male and hermaphrodite body size (measures in microns). Strain and sex had significant effects on body length (ANOVA, effect *sex*: F_1,422_=2325.80, P<0.0001; effect *strain:* F_10,422_=12.78, P<0.0001; interaction *sex* x *strain*: F_10,422_ =23.57, P<0.0001) and body width (ANOVA, effect *sex*: F_1,422_=1549,46, P<0.0001; effect *strain:* F_10,422_=15.25, P<0.0001; interaction *sex* x *strain*: F_10,422_ =19.47, P<0.0001). (ANOVA analyses testing for the fixed effects of *strain* and *sex* on body length/width; all data log-transformed).

| Strain | Sex | Body length | SE length | Body width | SE width | N individuals |
| --- | --- | --- | --- | --- | --- | --- |
| LSJ1 | M | 716.5 | 7.9 | 40.9 | 0.6 | 24 |
| LSJ1 | H | 998 | 11.2 | 62 | 0.9 | 23 |
| JU561 | M | 732.9 | 8.1 | 43.1 | 0.4 | 19 |
| JU561 | H | 1012 | 13.3 | 56 | 1.0 | 18 |
| CX11285 | M | 722.2 | 12.5 | 40.2 | 0.8 | 19 |
| CX11285 | H | 960 | 8.0 | 62 | 0.6 | 18 |
| N2 | M | 786.8 | 8.1 | 43.7 | 0.6 | 23 |
| N2 | H | 1057 | 9.2 | 62 | 0.7 | 20 |
| DL238 | M | 794.7 | 8.2 | 45.2 | 0.9 | 21 |
| DL238 | H | 994 | 12.0 | 63 | 1.1 | 17 |
| JU751 | M | 822.3 | 10.0 | 43.4 | 0.7 | 15 |
| JU751 | H | 1120 | 13.8 | 63 | 1.2 | 17 |
| JU1200 | M | 800.1 | 7.8 | 46.6 | 0.3 | 21 |
| JU1200 | H | 1140 | 13.2 | 67 | 0.8 | 21 |
| CB4856 | M | 747.8 | 7.9 | 43.6 | 0.4 | 21 |
| CB4856 | H | 965 | 8.3 | 60 | 0.4 | 19 |
| EG4946 | M | 716.1 | 12.1 | 47.2 | 1.2 | 10 |
| EG4946 | H | 974 | 9.1 | 65 | 1.1 | 16 |
| JU782 | M | 819.0 | 5.8 | 48.4 | 0.8 | 24 |
| JU782 | H | 972 | 10.7 | 62 | 0.9 | 15 |
| JU393 | M | 764.0 | 6.9 | 47.2 | 0.7 | 17 |
| JU393 | H | 1018 | 9.0 | 62 | 0.7 | 29 |

